# Assessment of long-term trends in genetic mean and variance after the introduction of genomic selection in layers: a simulation study

**DOI:** 10.1101/2023.02.20.529187

**Authors:** Ivan Pocrnic, Jana Obšteter, R. Chris Gaynor, Anna Wolc, Gregor Gorjanc

**Affiliations:** The Roslin Institute and Royal (Dick) School of Veterinary Studies, The University of Edinburgh, Edinburgh, United Kingdom; Agricultural Institute of Slovenia, Ljubljana, Slovenia; Iowa State University, Ames, IA, USA; Hy-Line International, Dallas Center, IA, USA

**Keywords:** genomic selection, stochastic simulation, optimal contributions, long-term selection, layers

## Abstract

Nucleus-based breeding programs are characterized by intense selection that results in high genetic gain, which inevitably means reduction of genetic variation in the breeding population. Therefore, genetic variation in such breeding systems is typically managed systematically, for example, by avoiding mating the closest relatives to limit progeny inbreeding. However, intense selection requires maximum effort to make such breeding programs sustainable in the long-term. The objective of this study was to use simulation to evaluate the long-term impact of genomic selection on genetic mean and variance in an intense layer chicken breeding program. We developed a large-scale stochastic simulation of an intense layer chicken breeding program to compare conventional truncation selection to genomic truncation selection optimized with either minimization of progeny inbreeding or full-scale optimal contribution selection. We compared the programs in terms of genetic mean, genic variance, conversion efficiency, rate of inbreeding, effective population size, and accuracy of selection. Our results confirmed that genomic truncation selection has immediate benefits compared to conventional truncation selection in all specified metrics. A simple minimization of progeny inbreeding after genomic truncation selection did not provide any significant improvements. Optimal contribution selection was successful in having better conversion efficiency and effective population size compared to genomic truncation selection, but it must be fine-tuned for balance between loss of genetic variance and genetic gain. In our simulation, we measured this balance using trigonometric penalty degrees between truncation selection and a balanced solution and concluded that the best results were between 45° and 65°. This balance is specific to the breeding program and depends on how much immediate genetic gain a breeding program may risk vs. save for the future. Furthermore, our results show that the persistence of accuracy is better with optimal contribution selection compared to truncation selection. In general, our results show that optimal contribution selection can ensure long-term success in intensive breeding programs using genomic selection.

## 1 Introduction

Genomic selection is a mature technology that is routinely applied in commercial animal and plant populations. It has a recognized positive effect on genetic gain, which in most applications exceeds conventional selection (Meuwissen et al., 2001; Schaeffer, 2006; Wiggans et al., 2017). Here, conventional selection is defined as the best linear unbiased prediction (BLUP) based on the pedigree relationship matrix (Henderson, 1984). In contrast, genomic selection is defined as genomic BLUP (GBLUP) based on a genomic relationship matrix constructed from the set of dense genome-wide single nucleotide polymorphism (SNP) markers (Meuwissen et al., 2001; VanRaden, 2008), or more often, a single-step GBLUP (ssGBLUP) based on a combination of genomic and pedigree relationship matrices (Legarra et al., 2009; Christensen and Lund, 2010). The superiority of genomic selection over conventional selection comes from its power to provide more accurate breeding values for young animals without their own phenotype, due to its ability to capture the Mendelian sampling, and consequently earlier selection leading to a decreased generation interval. Furthermore, genomic selection can enhance evaluations of difficult and expensive-to-measure traits as well as traits with low heritability (Calus et al., 2013).

While the combination of high accuracy for young animals and a short generation interval are the main drivers of increased genetic gain with genomic selection, the fuel for a successful selection process is the genetic variation in the population under selection. Inevitably, the theory states that during the selection process, genetic variance will decline due to changes in allele frequency caused by selection and random drift, and due to the accumulation of negative linkage disequilibrium, the Bulmer effect (Bulmer, 1971; Lynch et al., 1998; Walsh and Lynch, 2018). Hence, breeding programs monitor and manage genetic variation to avoid rapid reduction in effective population size (*Ne*) that can threaten the sustainability of future genetic gains. Interestingly, the trends of genetic mean reported in most conventional selection breeding programs are stable, suggesting room for future genetic gains even after intense conventional selection in the recent years or decades (for discussion, see Hill (2016)). The classical way to assess genetic variation in a population is to estimate the rate of inbreeding or equivalently *Ne*. There is a limited number of studies with retrospective analysis of genetic variance trends (Hidalgo et al., 2020) and even fewer studies properly dissecting the processes that drive these variance trends (Macedo et al., 2021; Lara et al., 2022). In the short-term, genomic selection is unquestionably demonstrating an increased rate of genetic gain per unit of time. Genomic selection also has the ability to reduce the rate of coancestry via more precise estimates of the Mendelian sampling terms between siblings and hence better control of future population and individual inbreeding (Daetwyler et al., 2007; Meuwissen et al., 2020). The studies examining the long-term effects of genomic selection are very scarce, e.g., see Gorjanc et al. (2018) and Wientjes et al. (2022). Furthermore, while we have traditionally described the genetic variation of populations using pedigree-based information, we can and should shift to more informative measures based on genomic information (Sonesson et al., 2012; Meuwissen et al., 2020).

Commercial layer chicken breeding programs are characterized by an intensive selection of elite purebred animals inside the closed lines. Genetic variation in such breeding systems is typically managed systematically, for example, by avoiding mating the closest relatives to limit progeny inbreeding. However, intense selection requires maximum effort to make layer breeding programs sustainable in the long-term. This could be achieved with optimal contribution selection (OCS), which maximizes genetic gain for a targeted rate of coancestry and, as such, manages future relationships between individuals in addition to progeny inbreeding (Woolliams et al., 1999). Technically, OCS optimizes genetic contributions of selection candidates to maximize a selection criterion (most commonly estimated breeding values) while constraining the group coancestry between these individuals. The mean of the selection criterion weighted by the optimised contributions is a measure of future genetic mean, while group coancestry weighted by the optimised contributions is a measure of future group coancestry. The advantage of OCS over truncation selection is its emphasis on managing between and within family (Mendelian sampling) variation (Howard et al., 2018). The usefulness of OCS in conventional layer breeding programs was presented by König et al. (2010). However, there is a lack of studies showing the long-term impact of genomic optimal contribution selection in intense layer breeding programs characterized by very short generation intervals and high selection intensity.

The objective of this study was to use simulation to evaluate the long-term impact of genomic selection on genetic mean and variance in an intense layer chicken breeding program. The simulation parameters were based on estimates from real data. Truncation genomic selection was compared to conventional selection and various OCS scenarios to fully explore the balance between maximizing genetic gain and managing genetic variation of a breeding program. Understanding and assessing this balance over a longer period provides a valuable decision-making platform for intense breeding programs with a focus on short-term competitiveness and long-term sustainability.

## 2 MATERIALS AND METHODS

We analyzed how different breeding scenarios impact the genetic mean and variance in an intense layer chicken breeding program over 30 years under selection. These scenarios included conventional and genomic truncation selection with random mating, genomic truncation selection with optimized mating to minimize progeny inbreeding, and two instances of genomic optimal contribution selection. Here, we first describe the stochastic simulation of a commercial layer breeding program according to the real parameters. Second, we provide details of the aforementioned breeding scenarios, including how we estimated breeding values and how we estimated optimal contribution selection. Finally, we describe the measures used to compare the scenarios (conversion efficiency, rate of inbreeding, *Ne*, and accuracy).

### 2.1 Stochastic simulation of a layer breeding program

We used the `AlphaSimR` package (Gaynor et al., 2021) to simulate 30 years of a commercial layer chicken line-breeding program. We initiated the simulation by generating base population genomes for 2,500 individuals using the Markovian coalescent simulator MaCS (Chen et al., 2009) as implemented in `AlphaSimR`. The simulated genomes mimicked the chicken genome with 39 autosomal chromosomes, total genetic length of 30 Morgans, and a total physical length of 1.2 × 10^9^ base pairs. To keep the simulation parameters consistent across all the chromosomes, we assumed they were all the same size. This departs from reality as the chicken genome consists of several microchromosomes. The recombination rate was set to 2.5 × 10^*−*8^ and the mutation rate to 5.0 × 10^*−*8^. While we used this recombination rate also in next simulation steps (see the next section), we have assumed that mutation is absent in the next simulation steps. This is an important caveat of this and similar simulation studies, as adding a realistic level of mutation to gene drop simulation of whole genomes is non-trivial. Efficient methods for simulating mutations are currently under development (Baumdicker et al., 2022). According to the demography of chickens, including domestication and selective breeding, the base population *Ne* was set to 100, with a gradual decrease from *Ne* of 500,000 at about 1 million generations ago. We retained 2,250 segregating sites per chromosome (87,750 in total) in the base population. Out of those, we selected at random 250 sites per chromosome (9,750 total) as quantitative trait loci (QTL), and 1,000 sites per chromosome (39,000 total) as single nucleotide polymorphism (SNP) markers to be used for genomic selection. Further 1,000 sites per chromosome (39,000 total) served as seemingly neutral loci used for monitoring genetic variation at loci not under direct selection. There was no overlap between QTL, SNPs, and neutral loci. To mimic the egg production phenotypes during the productive life of a hen, we simulated three purely additive traits for early, mid, and late egg production, having respective heritabilities of 0.18, 0.22, 0.25 with a correlation 0.75 between trait 1 and trait 2, 0.70 between trait 2 and trait 3, and 0.60 between trait 1 and trait 3 in the base population. The simulation and all the subsequent analyses were replicated 5 times to assess variability between the ‘biological’ replicates of the simulation.

### 2.2 Breeding scenarios

We evaluated five different breeding programs. Each breeding program started from a 10-year burn-in that used a conventional truncation selection on BLUP and random mating with equal contributions. Burn-in was followed by a 20-year evaluation period that used a: (i) continuation of the conventional truncation selection based on BLUP and random mating with equal contributions (PTS); (ii) genomic truncation selection based on ssGBLUP and random mating with equal contributions (GTS); (iii) genomic truncation selection based on ssGBLUP and optimized mating minimizing progeny inbreeding with equal contributions (GTSMF); (iv) Genomic optimal contribution selection based on ssGBLUP with a constrained number of sires and random pairing of the optimized contributions (GOCS); and (v) Genomic optimal contribution selection based on ssGBLUP with an unconstrained number of sires and random pairing of the optimized contributions (UGOCS). Additionally, we simulated a random selection program as a negative control to validate the *Ne* estimates.

A single year of the conventional or genomic breeding program is shown in Figure 1. In the programs, we mated 1,080 dams with either 40 or 120 sires. Therefore, the ratio of sires to dams was 1:27 for the 40 sires scenario and 1:9 for the 120 sires scenario. These values (9 and 27) were also absolute sire contributions under random mating and constrained optimized mating breeding programs. We assumed each dam had an equal contribution of her initial 9 female and 4 male offspring, which resulted in 9,720 female and 4,320 male selection candidates in each breeding cycle, 14,040 in total. One year of a conventional program allowed for one round of selection, and mating after female phenotypes were collected and used for genetic evaluation. In contrast, genomic programs reduced the age of sires by half, allowing two rounds of sire and dam selection per year, though dams were mated after one generation when their phenotypes became available. Selection was based on an index of breeding values for the three traits, with respective weights of 0.20, 0.35, and 0.45, thus giving more emphasis on the traits measured later in the lifetime as commonly used in breeding companies.

**Figure 1.**
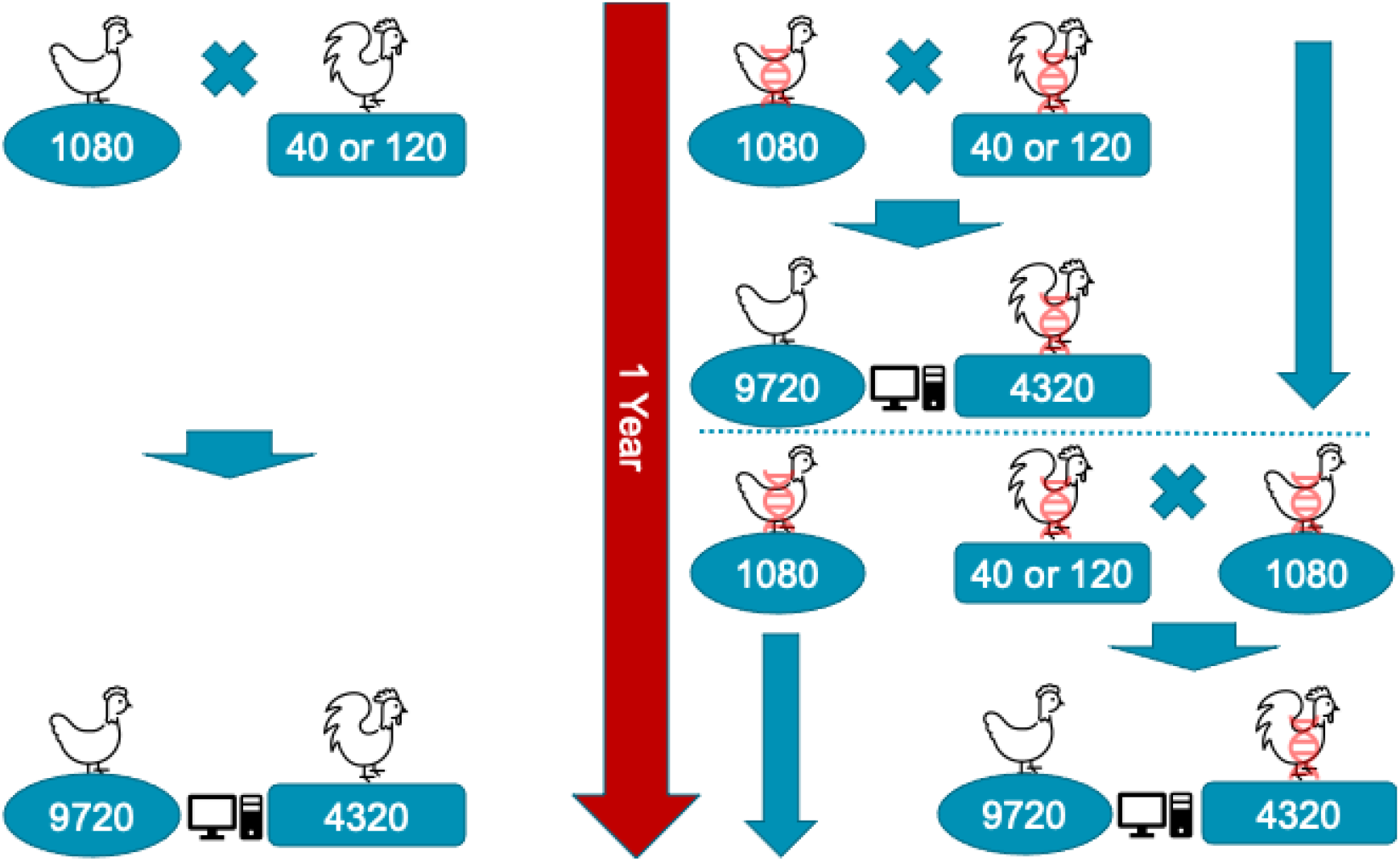
Schematic representation of a single year of conventional (left) and genomic breeding programs (right) with 1,080 dams mated with 40 or 120 sires generating 9,720 female and 4,320 male selection candidates. In the genomic programs animals marked with DNA helix were genotyped, which enabled earlier selection, though females were still mated after one generation causing generations to overlap.

In the truncation selection scenarios without optimization (PTS and GTS), we selected the top 1,080 females and either the top 40 or 120 males as the next-generation parents based on the index of breeding values obtained by either BLUP or ssGBLUP, and mated the parents at random with equal contributions. In the optimized scenarios, we used `AlphaMate` software (Gorjanc and Hickey, 2018) for a) optimized mating by minimizing future progeny inbreeding (GTSMF) and b) genomic optimal contribution selection with a targeted rate of coancestry with or without constraining the number of sires (GOCS and UGOCS).

In the GTSMF breeding program, we first selected the top 1080 females and either the top 40 or 120 males and then optimized their matings with regard to minimizing progeny inbreeding. For this optimisation we passed to `AlphaMate` an index of breeding values **a** obtained by ssGBLUP and a pedigree-based numerator relationship matrix **A** between the selected candidates. This optimization used an evolutionary algorithm that chose pairs of the selected candidates that minimised progeny inbreeding, that is, we minimized the coancestry between the pairs of parents.

In the GOCS breeding programs, we selected 1,080 females and 40 or 120 males that maximized the genetic gain under a targeted rate of coancestry. In the UGOCS breeding programs, we selected 1,080 females and any number of males that maximized the genetic gain under a targeted rate of coancestry. Additionally, we tested UGOCS starting from two different starting points; after the burn-in, using 40 or 120 sires. In the results, we report only the UGOCS that started after the burn-in with 120 sires. In UGOCS, we removed solutions that did not meet biological or logistical constraints in terms of the number of sires per generation. We set these limits between 20 and 200 sires per generation. We ran the optimisation only on the male side as optimising both male and female contributions was slow and required a lot of computing resources. Also, OCS can account for the previous selection (Henryon et al., 2015), here the selection of females when optimising male contributions. For these optimisations we passed to `AlphaMate` an index of breeding values **a** obtained by ssGBLUP and a pedigree-based numerator relationship matrix **A** between selection candidates. This optimisation used an evolutionary algorithm that choose optimal contributions of selection candidates. Specifically, the goal was to maximize **x**^*T*^ **a**, where **x** is a vector of contributions of selection candidates to the next generation [0, 0.5], while constraining the selected group coancestry 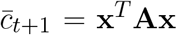 relative to current group coancestry 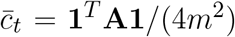, where *m* is number of matings, such that we obtained the targeted rate of coancestry 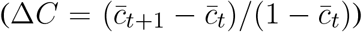 and the associated effective population size *Ne* = 1/(2∆*C*). We used the pedigree-based numerator relationship matrix, instead of the genome-based numerator relationship matrix, following the results from Meuwissen et al. (2020).

For GOCS and UGOCS, the balance between genetic gain and the rate of coancestry was optimized following Kinghorn (2011) with ‘trigonometric penalty degrees’ between the maximal genetic gain solution and the targeted solution under biological or logistic constraints (number of sires in this study). In that sense, the maximal genetic gain solution is obtained with a sole maximization of the genetic gain **x**^*T*^ **a** under biological or logistic constraints (without considering genetic diversity), and gives a trigonometric penalty degree of 0°. The minimal loss of genetic diversity is obtained by the minimization of selected group coancestry **x**^*T*^ **ax** under biological or logistic constraints (without considering the genetic gain), and gives a trigonometric penalty degree of 90°. Therefore, targeting trigonometric penalty degrees of 45° equalizes genetic gain and maintenance of genetic diversity, targeting trigonometric penalty degrees of 0° is equal to the truncation selection, and targeting trigonometric penalty degrees of 90° represents conservation programs. We have optimized across a wide range of trigonometric penalty degrees (5°-85°) and reported results for the selected trigonometric penalty degrees that facilitated comparison with other programs and discussion of their properties.

#### 2.2.1 Breeding value estimation

We estimated breeding values using `BLUPF90` (Misztal et al., 2018) by running pedigree-based BLUP for the conventional program (PTS) or ssGBLUP for genomic programs (GTS, GTSMF, GOCS, UGOCS). They all used a three-trait linear mixed model:

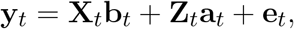

where **y**_*t*_ is a vector of phenotypes for the trait *t* (where *t* = *T* 1, *T* 2, *T* 3), **X**_*t*_ is a design matrix connecting the phenotype to mean as the only fixed effect **b**_*t*_, **Z**_*t*_ is a design matrix connecting the phenotypes to the animal breeding values **a**_*t*_, and **e**_*t*_ is a vector of residuals.

Variance components were assumed to be known using the base population simulation parameters, with their (co)variance structure being:

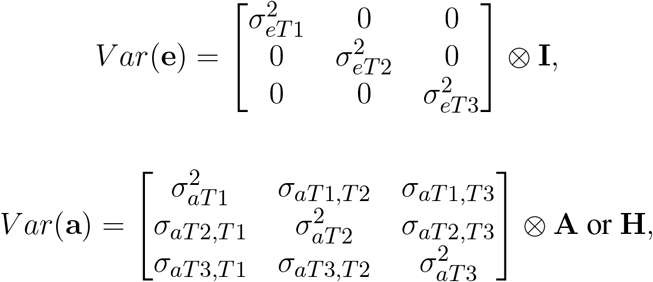

for BLUP and ssGBLUP, respectively, where **I** is the identity matrix, **A** is the pedigree-based numerator relationship matrix, and **H** is the matrix that combines pedigree and genomic relationships (Legarra et al., 2009; Christensen and Lund, 2010).

The **H** matrix was defined as:

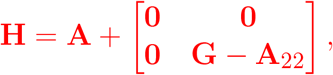

where **A**_22_ is pedigree-based numerator relationship matrix for genotyped animals only, 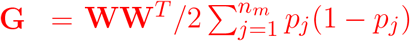, where **W** is a centred matrix of SNP genotypes (coded as 0 for the reference homozygote, 1 for the heterozygote, and 2 for the alternative homozygote), *p*_*j*_ is the observed frequency of the alternative allele for SNP *j*, and *n*_*m*_ is the number of SNPs (VanRaden, 2008).

For genomic evaluation, genotypes were available for 4,320 male selection candidates only and up to four generations back from previously selected males and females. Therefore, within each round of genomic selection, at most 8,800 and 9,120 genotypes were available respectively for 40 and 120 sires scenarios. The pedigree and phenotypic data were not truncated. The option `thrStopCorAG` was used to prevent `BLUPF90` from stopping when the correlations between **G** and **A**_**22**_ fell slightly below the default value (0.30). This option was used to overcome similar issues observed in some real datasets (e.g., Pocrnic et al. (2018)). All other `BLUPF90` settings were kept as default.

### 2.3 Comparison of breeding programs

We compared breeding scenarios in terms of genetic gain, genic standard deviation, conversion efficiency, rate of inbreeding, *Ne*, and accuracy of selection. To make the breeding scenarios comparable, all scenarios were normalized to the last year of burn-in (year 10), so that the mean genetic value was 0 and the standard deviation was 1. For each of the 20 evaluation years (11-30), we reported average values across 14,040 male and female selection candidates. In the genomic programs, we had two batches of selection candidates (28,080 total) per year, which we accounted for in the analyses by adding half a year points (e.g., 11, 11.5, 12, 12.5, …).

We measured genetic gain as the average true genetic value per year of birth (including half-year points in genomic programs). Genic standard deviation was calculated as the square root of the variance of true genetic values under the assumption of no linkage between the causal loci, that is 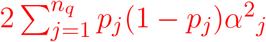 where *p*_*j*_ is the frequency of the alternative allele for QTL at locus *j, n*_*q*_ is the number of QTL, and *α*_*j*_ is the QTL additive effect at locus *j* (e.g., Lara et al. (2022)). The true genetic values and corresponding true variances were obtained directly from the AlphaSimR simulation. We measured the long-term viability of breeding programs through conversion efficiency (Gorjanc et al., 2018). The conversion efficiency was calculated by regressing the achieved genetic gain on the lost genic standard deviation. Within this definition, the conversion efficiency is the slope of the aforementioned linear regression, that is, the genetic gain that could be achieved when all the genic variance is utilized. Thus, the conversion efficiency can be useful for assessing the sustainability of a breeding program, as it combines measures of the gain and the diversity in a single metric, allowing easy extrapolation to the future, and informing the breeder how efficient is the breeding program in transforming the variance into gain. The average genomic inbreeding coefficients in year *t* (*F*_*t*_) were calculated from the observed heterozygosity as *F*_*t*_ = 1 − *Ho*_*t*_, where *Ho*_*t*_ is the average observed heterozygosity in year *t*. We separately calculated the observed heterozygosity for QTL, SNP marker loci used in genomic selection, and neutral loci. For comparison purposes, we also calculated individual pedigree-based inbreeding coefficients using the Meuwissen and Luo (1992) algorithm implemented in the `RENUMF90` software (Misztal et al., 2018). From the average inbreeding coefficients per year, we calculated the rate of inbreeding as ∆*F* = 1 − *exp*(*β*), where *β* is the regression coefficient obtained by regressing the natural logarithm of (1 − *F*_*t*_) on the year of birth *t* (Pe’
srez-Enciso, 1995). The *Ne* was calculated as *Ne* = 1/(2*L*∆*F*), where ∆*F* is the rate of inbreeding per year and *L* is the generation interval defined as the average age of the parents at the birth of their offspring. The generation interval was 1.00 for the conventional program and 0.75 for the genomic programs. Additionally, we estimated *Ne* following Wright (1931) classical formula as 4*N*_*m*_*N*_*f*_ /(*N*_*m*_ + *N*_*f*_), where *N*_*m*_ and *N*_*f*_ are respectively number of sires and dams. We measured the accuracy of selection as the correlation between the estimated breeding values and the true genetic values. Whenever the reported metrics were compared in the terms of percentage differences, we applied the percentage change formula as ((New value − Base value)/(Base value)) ∗ 100. For example, when assessing percentage change between GTS and PTS, GTS would be New value and PTS would be Base value.

## 3 RESULTS

### 3.1 Conversion efficiency

Table 1 shows the mean genetic gain and genic standard deviation from the last generation of selection candidates, and the conversion efficiency of breeding scenarios. To accompany the table, Figure 2 shows the conversion efficiency trends for the scenarios, together with extrapolation to 50% genic variance lost. Supplementary Figure 1 shows the extrapolation to 100% genic variance lost.

**Table 1.** Mean genetic gain and mean genic standard deviation (SD) for the last generation of selection candidates together with overall conversion efficiency (SD over replicates in parentheses).

| Breeding program | 40 sires per generation |  |  | 120 sires per generation |  |  |
| --- | --- | --- | --- | --- | --- | --- |
|  | Genetic gain | Genic SD | Conversion efficiency | Genetic gain | Genic SD | Conversion efficiency |
| PTS | 12.9 (0.9) | 0.57 (0.03) | 30.4 (1.6) | 14.2 (0.3) | 0.76 (0.02) | 50.1 (3.9) |
| GTS | 19.0 (0.7) | 0.58 (0.03) | 46.4 (3.4) | 20.7 (0.9) | 0.67 (0.03) | 62.3 (5.6) |
| GTSMF | 19.3 (1.3) | 0.57 (0.03) | 46.1 (3.9) | 20.3 (0.9) | 0.68 (0.02) | 61.1 (3.4) |
| GOCS 45 | 17.9 (1.2) | 0.69 (0.03) | 55.1 (6.5) | 17.9 (0.8) | 0.74 (0.01) | 66.9 (2.6) |
| GOCS 65 | 13.6 (0.9) | 0.79 (0.02) | 62.8 (5.9) | 13.0 (0.6) | 0.83 (0.01) | 76.0 (6.8) |
| Fluctuating number of sires per generation |  |  |  |  |  |  |
|  | Genetic gain | Genic SD | Conversion efficiency |  |  |  |
| UGOCS 45 | 21.2 (0.8) | 0.56 (0.02) | 47.0 (1.4) |  |  |  |
| UGOCS 55 | 19.7 (0.9) | 0.71 (0.01) | 65.6 (2.2) |  |  |  |

PTS - conventional truncation selection; GTS - genomic truncation selection;
GTSMF - GTS with minimization of progeny inbreeding;

**Figure 2.**
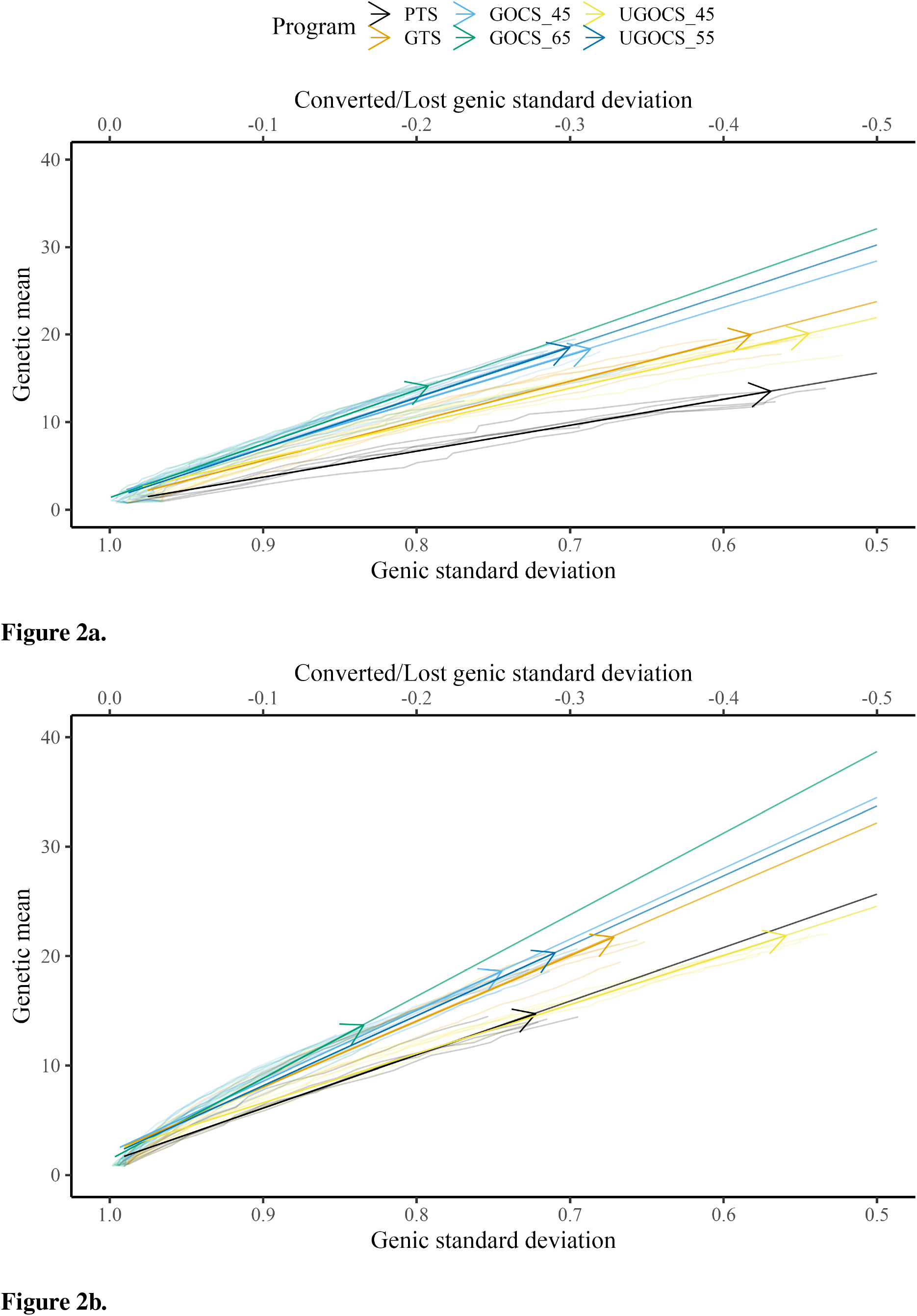
Conversion efficiency for conventional truncation selection (PTS) program and genomic programs (genomic truncation selection - GTS, genomic optimal contribution selection - GOCS X, unconstrained GOCS - UGOCS X, with the X trigonometric penalty degrees) marked with an arrow and further extrapolated to 50% of genic variance lost for **(A)** 40 sires and **(B)** 120 sires scenario.

The GTS delivered about 50% higher genetic gain than PTS; 19.0 vs. 12.9 genetic standard deviations in the 40 sires scenario, and 20.7 vs. 14.2 genetic standard deviations in the 120 sires scenario. Genic standard deviation was comparable between the PTS and GTS in the 40 sires scenario, about 0.58, giving a conversion efficiency of 30.4 for the PTS and 46.4 for the GTS. In the 120 sires scenario, both genetic gain and genic variance were larger than in the 40 sires scenario, giving a conversion efficiency of 50.1 for the PTS and 62.3 for the GTS.

The GOCS increased conversion efficiency by maximizing genetic gain at a targeted rate of coancestry. With a target of 45° trigonometric penalty degrees, GOCS had a somewhat similar genetic gain to the GTS, while simultaneously increasing conversion efficiency for 10% in the 40 sires scenario and for 7% in the 120 sires scenario. With a target of 65° trigonometric penalty degrees, GOCS had a similar genetic gain to the PTS, while simultaneously increasing conversion efficiency for 35% in the 40 sires scenario and for 22% in the 120 sires scenario. The GOCS had higher conversion efficiency in the 120 sires scenario compared to the 40 sires scenario.

The number of sires in UGOCS scenarios fluctuated as presented in the Supplementary Figure 2, with the general tendency that increasing the targeted trigonometric penalty degrees, that is, increasing emphasis on the maintenance of genetic variation, increased the number of sires. The average number of sires ranged from 2 for the target of 15° to 845 for the target of 85°. In the results, we focus on the UGOCS with a target of 45° and 55°, which respectively used 20 and 182 sires on average, and were therefore within the limits we set for the number of sires per generation (20-200). Furthermore, since the UGOCS starting from either of the two starting points (after 40 or 120 sires burn-in) tended to select a similar number of sires (Supplementary Figure 2), we report the results only for the one that started after the 120 sires burn-in. With a target of 45° trigonometric penalty degrees, UGOCS had a similar genetic gain, genic standard deviation, and conversion efficiency as the GTS. With a target of 55° trigonometric penalty degrees, UGOCS had a similar genic standard deviation and conversion efficiency of the GOCS with a target of 45° trigonometric penalty degrees.

### 3.2 Rate of inbreeding and effective population size

The rate of inbreeding per year and per generation, and the corresponding Ne in the 40 or 120 sires scenarios are presented in Table 2. We report rates of inbreeding multiplied by 100. Based on the average observed SNP heterozygosity, we obtained small estimates of *Ne* (*<* 40) in all breeding programs and scenarios. The GTS had on average somewhat larger *Ne* than the PTS. The GOCS scenarios with a target of 45 or 65 trigonometric penalty degrees had the largest *Ne*. For the 40 sires scenarios, the GOCS with a target of 65 trigonometric penalty degrees approximately doubled the *Ne* compared to the GTS (*Ne* 30 vs. 14) and approximately tripled the *Ne* compared to PTS (*Ne* 30 vs. 10). Similar increase was observed in the 120 sires scenarios. The rate of inbreeding and corresponding *Ne* values obtained by average observed SNP heterozygosity were very similar to the values obtained by measuring heterozygosity on neutral loci (Supplementary Table S1) or QTL (Supplementary Table S2). We expectedly observed larger *Ne* when using 120 sires compared to 40 sires. On average, using 120 sires instead of 40 sires increased the *Ne* by 37%, with the most considerable increase for PTS (60%). As expected, the UGOCS scenario with a target of 45° resulted in the lowest *Ne* (12), considering the low average number of sires (20). The UGOCS scenario with a target of 55° used on average 182 sires and resulted in *Ne* of 21, similar to GOCS scenarios with a target of 45° using either 40 or 120 sires (*Ne* of 19 and 23, respectively). Using the classical Wright’s formula, *Ne* was 154 and 432 respectively for the 40 and 120 sires scenarios. However, the above estimates of *Ne* from the observed rates of inbreeding depart significantly from the classical Wright’s formula due to intense selection over the 30 years. To validate our simulation and *Ne* estimates, we ran a random selection scenario. The random selection scenarios resulted in the *Ne* estimates of 151 for the 40 sires scenario and 517 for the 120 sire scenario, which were close to the classical Wright’s formula.

**Table 2.**
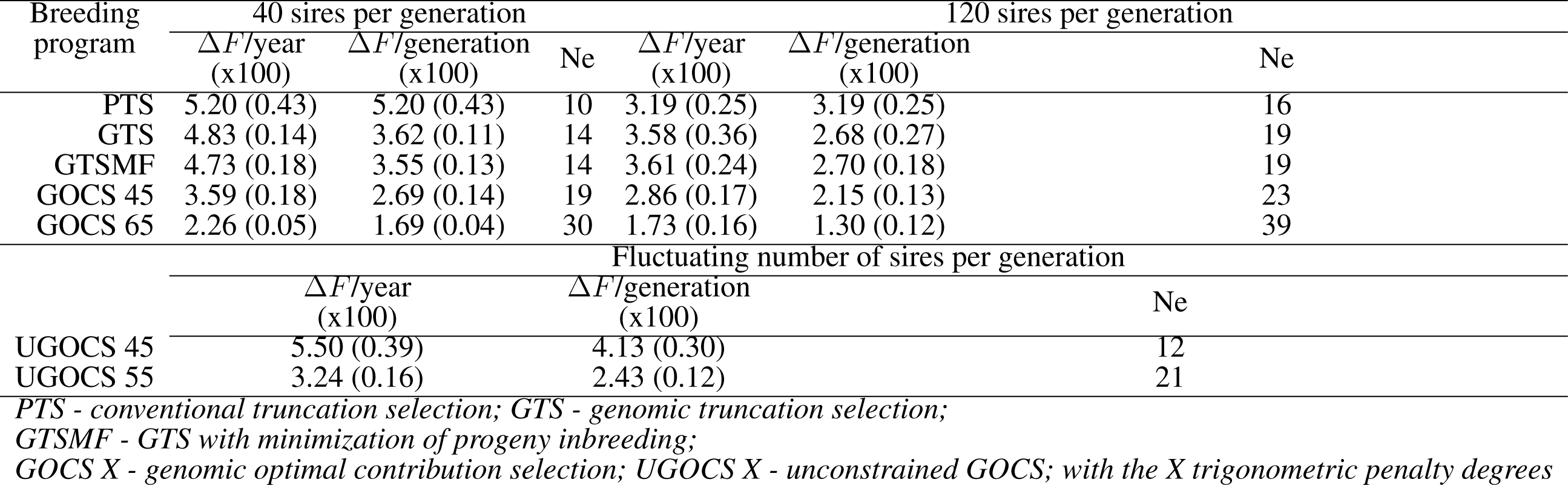
Rate of inbreeding (∆*F* x100) and effective population size (Ne) based on observed SNP heterozygosity (SD x100 over replicates in parentheses).

### 3.3 Accuracy of selection

The mean accuracy of selection over the 20 years is shown in Table 3, whereas the trends over the 20 years of selection are shown in the Supplementary Figure 3. The GTS had higher accuracy than PTS. The genomic data provided a 24% (40 sires) and 21% (120 sires) increase in overall accuracy compared to the conventional program based on phenotypic and pedigree data only. Compared to truncation selection programs, GOCS programs further increased accuracy. GOCS with a target of 65 trigonometric penalty degrees had the highest overall accuracy; 0.74 for the 40 sires scenario and 0.75 for the 120 sires scenario.

**Table 3.** Mean accuracy of selection candidates over 20 years of selection (SD over replicates in parentheses).

| Breeding program | 40 sires per generation | 120 sires per generation |
| --- | --- | --- |
| PTS | 0.50 (0.02) | 0.53 (0.01) |
| GTS | 0.64 (0.01) | 0.64 (0.01) |
| GTSMF | 0.64 (0.01) | 0.63 (0.01) |
| GOCS 45 | 0.69 (0.01) | 0.72 (0.01) |
| GOCS 65 | 0.74 (0.01) | 0.75 (0.01) |
| Fluctuating number of sires per generation |  |  |
| UGOCS 45 | 0.64 (0.01) |  |
| UGOCS 55 | 0.66 (0.01) |  |

*PTS* - conventional truncation selection; *GTS* - genomic truncation selection;
*GTSMF* - *GTS* with minimization of progeny inbreeding;
*GOCS X* - genomic optimal contribution selection; *UGOCS X* - unconstrained *GOCS*; with the *X* trigonometric p

Translated to the percentage increase, using GOCS with a target of 65 trigonometric penalty degrees increased the overall accuracy by 19% (40 sires) and 17% (120 sires) compared to the genomic truncation selection. On the other hand, there was no apparent advantage in using the UGOCS compared to the GTS. Furthermore, there were no significant differences in accuracy between using 40 or 120 sires. Looking at the accuracy trends across years (Supplementary Figure 3), the GOCS maintained accuracy the most. This beneficial trend was especially apparent in the initial evaluation years.

## 4 DISCUSSION

In this study, we affirmed that nucleus-based breeding programs, as used in commercial layer chicken breeding, successfully generated genetic gain with conventional selection and demonstrated a suitable structure to adopt genomic selection. Switching from the PTS to GTS increased genetic gain, predominantly through shortened generation intervals, and increased accuracy of selection for young animals. However, the nature of nucleus-based breeding programs requires managing genetic diversity to facilitate long-term genetic gain. Rapid genetic turnover with genomic selection makes this evergreen point even more important. Here, we developed and deployed a large-scale stochastic simulation of an intense layer chicken breeding program and evaluated the long-term impact of genomic selection on genetic mean and variance, and the effect of GOCS on efficiency. For this, we have simulated 30 years of breeding, and used last 20 years to compare the scenarios, which was considered a long-term period. Here, we discuss how GTS and GOCS affect: i) the long-term success of tested scenarios measured with the conversion efficiency; ii) the rate of inbreeding and *Ne*; and iii) the accuracy of selection.

### 4.1 Conversion efficiency

The GTS achieved 50% higher genetic gain than PTS with similar or even lower loss of genic variance. Genomic selection enables the within-family selection of genotyped selection candidates by estimating their parent average along with the Mendelian sampling term, compared to having only their parent average in conventional selection. Furthermore, genomic selection has decreased the generation interval and increased the accuracy of selection for young animals, as observed across livestock species, including layer chickens (Sitzenstock et al., 2013; Wolc et al., 2015; Picard Druet et al., 2020). For example, a two-to three-fold increase in response to selection was observed for egg production and quality traits with genomic compared to conventional selection (Sitzenstock et al., 2013; Wolc et al., 2015).

Besides genetic gain, we evaluated the sustainability of tested scenarios using conversion efficiency. The application of OCS is often seen as a risk-taking action that sacrifices some short-term genetic gain for long-term sustainability. Alternatively, we can assess the sustainability of the breeding program by looking at the annual trends of genetic gain and genic variance as presented respectively in Supplementary Figures 4 and 5. By looking at those trends, similarly to the conversion efficiency trends, we can sense how much short-term gain is sacrificed for generating more long-term gain with OCS. One of the key challenges in implementing OCS is choosing a balance between the selection and the management of genetic variation. In this study we follow the approach of Kinghorn (2011) who uses the ‘operational’ trigonometric penalty degrees between the truncation selection solution and the targeted optimal contribution selection solution (Gorjanc et al., 2018). In general, our results expectedly show that higher trigonometric penalty degrees result in larger conversion efficiency and lower short-term gain, while lower trigonometric penalty degrees result in genetic gain comparable to truncation selection. We have ran simulation with a wide range of trigonometric penalty degrees (5°-85°), but report only a subset of results that facilitated comparison between optimal and truncation selection scenarios. While we can run stochastic simulations across the grid of various trigonometric penalty degrees to find the best compromise for the desired goal, this might not be feasible in practical situations. Woolliams et al. (2015) suggest the target rate of coancestry should be less than 0.01 (*Ne >* 50), which can be a more concrete constraint in optimization than the trigonometric penalty degrees, which can vary the optimisation targets as input data and constraints change. In the results, we highlighted OCS scenarios with targets of 45° and 65° to demonstrate their value relative to truncation selection scenarios. The GOCS scenario with a target of 65° increased the conversion efficiency between 22% and 35%, and was hence the best strategy to achieve long-term sustainability at the expense of on average 33% lower short-term gain. Similarly, the GOCS scenario with a target of 45° increased conversion efficiency between 7% and 19%, which is useful for achieving short-term genetic gain while still investing in long-term sustainability. The results also indicate that the increase in the conversion efficiency of GOCS compared to GTS is a function of the number of breeding individuals since the increase was larger when using 40 sires than 120 sires. It is vital to point that for the same loss of genic variance both GOCS scenarios achieved higher genetic gain and conversion efficiency compared to GTS (Figure 2 and Supplementary Figure 1), which is the correct comparison between these scenarios. The reported optimization targets are specific to this study with its specific breeding program design and species-specific biology. For example, in a simulation study, Obšteter et al. (2019) reported the highest genetic gain for targets between 45° and 50° but with a conversion efficiency similar to their GTS, while their targets between 55° and 75° had much better conversion efficiency, but lower genetic gain compared to their GTS. Similarly, in our study, GOCS with targets higher than 65° achieved a lower genetic gain, even compared to conventional truncation selection (results not reported). We did not report results for targets lower than 45° as they had a similar gain and efficiency as the GTS. König et al. (2010) demonstrated the benefits of OCS compared to truncation selection in two commercial White Leghorn lines and one experimental line in the context of conventional selection. The advantage of OCS over truncation selection is in optimizing contributions, which are a function of animal Mendelian sampling terms (Woolliams et al., 2015; Howard et al., 2018). This means that OCS and genomic selection work in synergy (Daetwyler et al., 2007; Maltecca et al., 2020; Sonesson et al., 2012; Obšteter et al., 2019).

An additional benefit of employing GOCS is the potential optimization of the size of the breeding population. In our study, GOCS with a target of 65 trigonometric penalty degrees using 40 sires per generation resulted in similar conversion efficiency as GTS using 120 sires. Therefore, these results suggest that we can achieve the same long-term genetic gain, but not short-term genetic gain, with three times fewer sires per generation, which could reduce the production cost. UGOCS scenarios with a target of 45° and 55° resulted in a genetic gain comparable to GTS and, on average, selected 20 and 182 sires, which was biologically and logistically feasible. The UGOCS with a target of 55° used on average 182 sires and resulted in a conversion efficiency similar to that of GOCS with a target of 45° using 40 sires. Therefore, none of the UGOCS scenarios surpassed the short-term genetic gain of the GTS or achieved better long-term conversion efficiency compared to the GOCS scenarios. While there was no benefit in using the UGOCS, it was useful as a guideline to show that using less than 40 or more than 120 breeding males is not necessarily beneficial in our simulated breeding program. This is in line with a simulation of a pig breeding program in (Henryon et al., 2015) that has compared unconstrained and several constrained OCS scenarios. They concluded that the constrained scenarios achieved a similar long-term genetic gain compared to unconstrained scenarios.

### 4.2 Rate of inbreeding and effective population size

Although the conversion efficiency indicates the sustainability of a breeding program with respect to the traits under selection, the rate of inbreeding and associated *Ne* indicate broader sustainability, also including neutral diversity that encompasses potential future mutations, traits, or environments. We show that GTS improved the rate of inbreeding per generation compared to PTS, resulting in a larger *Ne*. This benefit of GTS comes from the ability to estimate Mendelian sampling terms, leading to better within-family differentiation and thus reducing the pressure on the co-selection of sibs from families with high parent averages in comparison to PTS (Daetwyler et al., 2007). In such comparisons, it is essential to measure the rate of inbreeding per generation rather than per year, because genomes are transmitted between generations. Not accounting for generation length in the comparison of the rate of inbreeding can lead to different conclusions, as it can be seen from the following example in scenarios using 120 sires. In this example, the rate of inbreeding per year with the GTS (∆*F*_*year*_ ∗ 100 = 3.58) is larger than with the PTS (∆*F*_*year*_ ∗ 100 = 3.19), while when measured per generation is lower (∆*F*_*generation*_ ∗ 100 = 2.68). On the other hand, when using 40 sires, the rate of inbreeding with GTS per generation (∆*F*_*generation*_ ∗ 100 = 3.62) as well as per year (∆*F*_*year*_ ∗ 100 = 4.83) was lower compared to PTS (∆*F*_*year*_ ∗ 100 = 5.20). These results are in line with Wolc et al. (2015), who used simulation of a layer breeding program and showed that GTS halved the rate of inbreeding per generation compared to PTS while keeping a similar inbreeding rate per year. Our *Ne* estimates were low, but not unexpected given the parameters of the simulation. We have simulated 30 years of intense selection with the rate of inbreeding (100) per generation between5.20 and 1.30 respectively giving *Ne* between 10 and 39. The estimates of *Ne* in livestock, including chickens, vary a lot across the literature due to intrinsic differences between populations, but also due to different estimation methods using pedigree or genomic data, different type of genomic data, different summaries of the data, different time points, and different types of *Ne*, for example see Waples (2022). Zhang et al. (2020) reported a *Ne* of 31 for Cornish, 109 for White Leghorn, and 189 for Rhode Island Red chicken breeds. For nine commercial pure lines with origin in Plymouth Rock and Cornish, Andreescu et al. (2007) reported *Ne* ranging from 50 to 200. In the study on two experimental (White Leghorn and New Hampshire pure lines) and two commercial (White Leghorn pure line and two-way cross between Rhode Island Red and White Rock) egg-layer lines, Qanbari et al. (2010) reported *Ne* to be less than 70 for brown layers and less than 50 for white layers. In the analysis of the Russian White breed, the *Ne* ranged from 14 to 124 (Dementieva et al., 2021). Pocrnic et al. (2016) approximated the *Ne* of 44 using a large commercial broiler chicken dataset. Compared to these estimates, our estimates are at a lower bound.

One possible reason for the discrepancy between our *Ne* estimates and those published could be our coalescent simulation of base population genomes, which is simulating neutral variation albeit at increasingly smaller *Ne* to mimic drift and selection due to domestication and recent selective breeding. Such simulations generate variation that has an abundance of rare variants with a typical U-shaped allele frequency spectrum (Daetwyler et al., 2013). While such variation can be captured by whole-genome sequencing in real populations, SNP arrays largely do not tag rare variants, and due to the SNP ascertainment bias they have uniform allele frequency spectrum, which can lead to mismatch between our and published (real data) estimates of *Ne*. This is also most likely why there were no major differences between the estimates of *Ne* obtained by average observed SNP heterozygosity and the estimates based on measuring heterozygosity on neutral loci or QTL. From the perspective of simulations, this might indicate that a good practice for future simulation studies is to consider sampling the SNPs with ascertainment bias. Still, we estimated the rate of inbreeding based on the rate of change in the observed heterozygosity, which was at the level of heterozygosity found in commercial chicken lines. Our simulations started in year 1 with observed marker heterozygosity (SD) of 0.25 (0.01), while our evaluated breeding scenarios (year 11; after the burn-in) started with observed heterozygosity of 0.16 (0.01) and 0.21 (0.01) respectively for 40 and 120 sires scenarios. These values matched values reported in the real data studies. Zhang et al. (2020) reported observed average heterozygosity for commercial lines ranging from 0.29 (0.02) to 0.39 (0.04), Qanbari et al. (2010) from 0.34 (0.15) to 0.47 (0.21), Elferink et al. (2012) from 0.21 to 0.43, Dementieva et al. (2021) from 0.31 to 0.39, and Malomane et al. (2019) from 0.12 to 0.28.

Minimizing progeny inbreeding after genomic truncation selection (GTSMF, a breeding strategy commonly used in practice) did not significantly affect genetic gain, genic variance, or conversion efficiency. Furthermore, this method resulted in the rates of inbreeding and corresponding *Ne*’s comparable to GTS, and therefore does not offer long-term advantage beyond short-term avoidance of progeny inbreeding and associated inbreeding depression. While inbreeding and associated inbreeding depression are important, for long-term sustainability managing coancestry between selected individuals and with this genetic variance is more important. In our study, GOCS with a target of 65 trigonometric penalty degrees provided the best properties for controlling the rate of inbreeding for both 40 and 120 sires scenarios. It approximately halved the rate of inbreeding and doubled the *Ne* compared to GTS, and approximately tripled the *Ne* compared to PTS. On the other hand, the UGOCS scenarios were unable to provide meaningful benefits for the rate of inbreeding compared to GOCS scenarios. In this study, UGOCS scenarios with a target of 45 and 55 trigonometric penalty degree used respectively on average 20 and 182 breeding sires. Therefore, a higher rate of inbreeding of UGOCS scenario with a target of 45° could be directly attributed to a low number of sires in each generation. In the UGOCS scenario with a target of 55°, the resulting rate of inbreeding was comparable to the GOCS scenario with a target of 45° using 40 sires. This reinforces the results we obtained for conversion efficiency, pointing out that having fewer than 40 sires seems risky, while having more than 120 sires does not provide any further benefit and only raises the costs of keeping more breeding individuals. König et al. (2010) evaluated OCS applied to the conventional breeding scheme of two commercial White Leghorn lines and one experimental line, and concluded that OCS is a preferred method for managing inbreeding in layer populations, supporting our results. Howard et al. (2018) studied closed nucleus commercial pig lines and connected the selective advantage of OCS to estimates of Mendelian sampling terms. Similarly to our study, they concluded that a combination of genomic selection and OCS has the potential to generate greater long-term genetic gain without a negative impact on the rate of inbreeding.

While genomic information is now the de-facto standard for estimating and managing genetic diversity (Baes et al., 2019; Howard et al., 2017), most studies also report pedigree-based inbreeding for comparison. The rationale for this comparison is that while pedigree data provide expected trends in genetic diversity over time, genome data provide actual genetic variation and hence realized trends in genetic diversity over time. However, caution is required in comparing pedigree- and genome-based estimates, because they might not be estimating the same quantity of interest or might be capturing different genetic processes driving the changes. In our simulation, we have observed large differences between the rates of inbreeding estimated from pedigree or genomic data. The pedigree-based estimates are presented in Supplementary Table S3. For example, in some scenarios, pedigree-based rates of inbreeding were about five to ten times smaller than genome-based rates of inbreeding, consecutively resulting in a tenfold larger estimate of *Ne*. In the scenarios using GOCS, estimates of the *Ne* from the pedigree-based inbreeding were even outside the range of estimates obtained by the random selection or Wright’s formula. The discrepancy between pedigree- and genome-based estimates is likely due to the fact that pedigree relationships model the expected drift and inbreeding under the infinitesimal model without actually observing changes in allele frequency and heterozygosity. With genomic data, we can observe such changes, which are also influenced by selection, and cannot be captured by pedigree data alone. In general, our genome-based estimates of rates of inbreeding exceeded the pedigree-based estimates across all the scenarios. This is in agreement with the existing literature across the livestock sector, noting that most of the reported genome-based estimates are based either on the genomic relationship matrix or runs of homozygosity. For example, in a study of Holstein and Jersey cattle populations, Makanjuola et al. (2020) reported ∆*F*_*generation*_ of 0.75 and 1.16 for pedigree-based and genome-based (runs of homozygosity).

The discrepancy between the pedigree- and genome-based inbreeding estimates (e.g., Table 2 and Supplementary Table S3) opens a much broader discussion on the proper management of genetic diversity in the genomics era. While in the conventional selection, both the evaluation and the optimal contribution steps typically use the same pedigree-based relationship matrix, this need not be the case with genomic selection. There are reports of the benefits of using various genome-based relationship matrices for optimal contribution selection (Gebregiwergis et al., 2020; Maltecca et al., 2020; Henryon et al., 2019; Sonesson et al., 2012; Woolliams et al., 2015; Meuwissen et al., 2020). In this study, we did not evaluate the impact of various relationship matrices to manage diversity in OCS, since only male selection candidates were genotyped in the simulation, mimicking the real-life breeding program. However, with the decreasing costs of genotyping, genotypes for young female selection candidates will eventually become available, allowing testing a full range of solutions to model relationships in the OCS, for example, see Meuwissen et al. (2020).

### 4.3 Accuracy of selection

Accuracy of selection is an essential metric in every breeding program, as it directly affects the amount of genetic gain that can be achieved. In our study, the GTS increased the average accuracy for the selection candidates by more than 20% compared to the PTS. This confirms the now well-established positive effect of genomic selection on the accuracy observed across livestock species, including chicken (Wolc et al., 2011; Sitzenstock et al., 2013; Picard Druet et al., 2020; Hidalgo et al., 2021). Despite the higher accuracy with genomic selection, its persistence in the long-term still has not been fully assessed (for example over 7 years, see Hidalgo et al. (2021)), especially when comparing truncation and optimal contribution selection. The rapid decline of the accuracy with GTS was shown by Muir (2007). They attributed the decline mainly to the erosion of the favorable linkage disequilibrium and showed that persistence is dependent on the size of the training population. Wolc et al. (2011) investigated the persistence of the accuracy using real data in layers and suggested that the decrease in the accuracy could be balanced by increasing the size of the training population (by accumulating data) and retraining the reference in every generation. Furthermore, the accuracy might also decrease over generations due to a drop in the genetic variance, which provides the opportunity for OCS. Hidalgo et al. (2021) reported that in general, accumulating data increased the accuracy but also that keeping only two most recent years of data (pedigree, phenotypes and genotypes) was sufficient to have persistent accuracies for selection candidates.

While we report the average accuracy of all selection candidates, it is worth mentioning that there will be a difference between the accuracy for male and female selection candidates due to the difference in available information in our breeding program. For example, in the last generation of selection in the 40 sires GTS scenarios, the average accuracy (SD) for male selection candidates was 0.70 (0.02) and for female selection candidates 0.46 (0.04). In GOCS with target of 45 and 65 trigonometric penalty degrees, the average accuracy for male selection candidates was respectively 0.76 (0.02) and 0.81 (0.03), and for female selection candidates was respectively 0.57 (0.04) and 0.66 (0.07).

We found that the average accuracy over 20 years increased by about 20% when we used the GOCS compared to GTS. Furthermore, the accuracy trends over the years were more stable when we used the GOCS. A similar boost in accuracy with OCS was reported in several studies (Gourdine et al., 2012; Eynard et al., 2018). In a simulation of a plant-breeding program with rapid recurrent genomic selection, Gorjanc et al. (2018) argued that the positive impact of OCS on accuracy is due to better management of genetic variation, so the genetic drift between training and prediction populations is not too large. (Jannink, 2010) showed that increasing the rate of inbreeding with genomic selection decreases the accuracy, indicating that managing the rate of inbreeding is beneficial for accuracy. Other methods of selection and mating optimization have shown a similar positive trend on the accuracy of selection, like the recently proposed scoping method in plant breeding (Vanavermaete et al., 2020). Although the underlying mechanism that drives the boost and persistence of the accuracy in the OCS is not yet fully understood, our results suggest the benefit of using it compared to truncation selection.

## 5 CONCLUSIONS

We developed a large-scale simulation of an intense layer chicken breeding program and evaluated the long-term impact of genomic selection on genetic gain and variance. With the transition from conventional to genomic truncation selection, we decreased the generation interval and increased the accuracy of selection for young animals. This resulted in a substantial increase in genetic gain with a similar loss of variance, meaning that genomic selection converted genetic variation to gain better compared to the conventional truncation selection. While the GOCS delivered somewhat lower genetic gain in the short-term compared to GTS, it managed genetic variance better and maintained high selection accuracy for longer. By optimizing the balance between the loss of genetic variance and genetic gain, GOCS had a better conversion efficiency and therefore showed the potential for larger long-term genetic gain compared to GTS. Furthermore, our results suggest that the application of GOCS can be useful to manage the number of parents. In general, our results indicate that the intense layer chicken breeding programs should use genomic selection and consider improving the conversion of genetic variation to gain with GOCS to ensure long-term success of their breeding programs. Consequently, the GOCS should be tested and deployed in all closed-nucleus breeding programs.

## AUTHOR CONTRIBUTIONS

IP co-designed the study, analyzed the data, interpreted the results, and wrote the manuscript. RCG contributed to simulation development and results interpretation. AW contributed layer breeding information and contributed to the interpretation of results. JO contributed to the interpretation of results and manuscript revision. GG initiated, co-designed, and supervised the study and contributed to all its stages. All authors reviewed and approved the manuscript.

## ACKNOWLEDGMENTS

We acknowledge John Woolliams (The University of Edinburgh, UK) for helpful discussions and members of the HighlanderLab on initial draft of the manuscript. This work has made use of the resources provided by the University of Edinburgh Compute and Data Facility (ECDF) (http://www.ecdf.ed.ac.uk).

## SUPPLEMENTAL DATA

Supplemetal tables and figures are in “frontiers SupplementaryMaterial.tex”

## DATA AVAILABILITY STATEMENT

R code for data simulation is publicly available at https://github.com/HighlanderLab/ipocrnic_Layer_OCS.

## REFERENCES

Andreescu, C., Avendano, S., Brown, S. R., Hassen, A., Lamont, S. J., and Dekkers, J. C. (2007). Linkage disequilibrium in related breeding lines of chickens. Genetics 177, 2161–2169

Baes, C. F., Makanjuola, B. O., Miglior, F., Marras, G., Howard, J. T., Fleming, A., et al. (2019). Symposium review: The genomic architecture of inbreeding: How homozygosity affects health and performance. Journal of dairy science 102, 2807–2817

Baumdicker, F., Bisschop, G., Goldstein, D., Gower, G., Ragsdale, A. P., Tsambos, G., et al. (2022). Efficient ancestry and mutation simulation with msprime 1.0. Genetics 220, iyab229

Bulmer, M. (1971). The effect of selection on genetic variability. The American Naturalist 105, 201–211

Calus, M., Berry, D., Banos, G., De Haas, Y., and Veerkamp, R. (2013). Genomic selection: the option for new robustness traits? Advances in Animal Biosciences 4, 618–625

Chen, G. K., Marjoram, P., and Wall, J. D. (2009). Fast and flexible simulation of dna sequence data. Genome research 19, 136–142

Christensen, O. F. and Lund, M. S. (2010). Genomic prediction when some animals are not genotyped. Genetics Selection Evolution 42, 1–8

Daetwyler, H. D., Calus, M. P. L., Pong-Wong, R., de los Campos, G., and Hickey, J. M. (2013). Genomic Prediction in Animals and Plants: Simulation of Data, Validation, Reporting, and Benchmarking. Genetics 193, 347–365. doi:10.1534/genetics.112.147983

Daetwyler, H. D., Villanueva, B., Bijma, P., and Woolliams, J. A. (2007). Inbreeding in genome-wide selection. Journal of Animal Breeding and Genetics 124, 369–376

Dementieva, N. V., Mitrofanova, O. V., Dysin, A. P., Kudinov, A. A., Stanishevskaya, O. I., Larkina, T. A., et al. (2021). Assessing the effects of rare alleles and linkage disequilibrium on estimates of genetic diversity in the chicken populations. Animal 15, 100171

Elferink, M. G., Megens, H.-J., Vereijken, A., Hu, X., Crooijmans, R. P., and Groenen, M. A. (2012). Signatures of selection in the genomes of commercial and non-commercial chicken breeds. PLoS One 7, e32720

Eynard, S. E., Croiseau, P., Lalo, D., Fritz, S., Calus, M. P., and Restoux, G. (2018). Which individuals to choose to update the reference population? minimizing the loss of genetic diversity in animal genomic selection programs. G3: Genes, Genomes, Genetics 8, 113–121

Gaynor, R. C., Gorjanc, G., and Hickey, J. M. (2021). Alphasimr: an r package for breeding program simulations. G3 11, jkaa017

Gebregiwergis, G. T., Sørensen, A. C., Henryon, M., and Meuwissen, T. (2020). Controlling coancestry and thereby future inbreeding by optimum-contribution selection using alternative genomic-relationship matrices. Frontiers in Genetics 11, 345

Gorjanc, G., Gaynor, R. C., and Hickey, J. M. (2018). Optimal cross selection for long-term genetic gain in two-part programs with rapid recurrent genomic selection. Theoretical and applied genetics 131, 1953–1966

Gorjanc, G. and Hickey, J. M. (2018). Alphamate: a program for optimizing selection, maintenance of diversity and mate allocation in breeding programs. Bioinformatics 34, 3408–3411

Gourdine, J.-L., Sørensen, A., and Rydhmer, L. (2012). There is room for selection in a small local pig breed when using optimum contribution selection: a simulation study. Journal of Animal Science 90, 76–84

Henderson, C. R. (1984). Applications of Linear Models in Animal Breeding (Guelph, Canada: University of Guelph)

Henryon, M., Liu, H., Berg, P., Su, G., Nielsen, H. M., Gebregiwergis, G. T., et al. (2019). Pedigree relationships to control inbreeding in optimum-contribution selection realise more genetic gain than genomic relationships. Genetics Selection Evolution 51, 1–12

Henryon, M., Ostersen, T., Ask, B., Sørensen, A. C., and Berg, P. (2015). Most of the long-term genetic gain from optimum-contribution selection can be realised with restrictions imposed during optimisation. Genetics Selection Evolution 47, 1–14

Hidalgo, J., Lourenco, D., Tsuruta, S., Masuda, Y., Breen, V., Hawken, R., et al. (2021). Investigating the persistence of accuracy of genomic predictions over time in broilers. Journal of Animal Science 99. doi:10.1093/jas/skab239. Skab239

Hidalgo, J., Tsuruta, S., Lourenco, D., Masuda, Y., Huang, Y., Gray, K. A., et al. (2020). Changes in genetic parameters for fitness and growth traits in pigs under genomic selection. Journal of animal science 98, skaa032

Hill, W. G. (2016). Is continued genetic improvement of livestock sustainable? Genetics 202, 877–881

Howard, D. M., Pong-Wong, R., Knap, P. W., Kremer, V. D., and Woolliams, J. A. (2018). Selective advantage of implementing optimal contributions selection and timescales for the convergence of long-term genetic contributions. Genetics Selection Evolution 50, 1–10

Howard, J. T., Pryce, J. E., Baes, C., and Maltecca, C. (2017). Invited review: Inbreeding in the genomics era: Inbreeding, inbreeding depression, and management of genomic variability. Journal of dairy science 100, 6009–6024

Jannink, J.-L. (2010). Dynamics of long-term genomic selection. Genetics Selection Evolution 42, 1–11

Kinghorn, B. P. (2011). An algorithm for efficient constrained mate selection. Genetics Selection Evolution 43, 1–9

König, S., Tsehay, F., Sitzenstock, F., Von Borstel, U., Schmutz, M., Preisinger, R., et al. (2010). Evaluation of inbreeding in laying hens by applying optimum genetic contribution and gene flow theory. Poultry Science 89, 658–667

Lara, L. A. d. C., Pocrnic, I., Oliveira, T. d. P., Gaynor, R. C., and Gorjanc, G. (2022). Temporal and genomic analysis of additive genetic variance in breeding programmes. Heredity 128, 21–32

Legarra, A., Aguilar, I., and Misztal, I. (2009). A relationship matrix including full pedigree and genomic information. Journal of dairy science 92, 4656–4663

Lynch, M., Walsh, B., et al. (1998). Genetics and analysis of quantitative traits, vol. 1 (Sinauer Sunderland, MA)

Macedo, F. L., Christensen, O. F., and Legarra, A. (2021). Selection and drift reduce genetic variation for milk yield in manech tete rousse dairy sheep. JDS communications 2, 31–34

Makanjuola, B. O., Miglior, F., Abdalla, E. A., Maltecca, C., Schenkel, F. S., and Baes, C. F. (2020). Effect of genomic selection on rate of inbreeding and coancestry and effective population size of holstein and jersey cattle populations. Journal of dairy science 103, 5183–5199

Malomane, D. K., Simianer, H., Weigend, A., Reimer, C., Schmitt, A. O., and Weigend, S. (2019). The synbreed chicken diversity panel: a global resource to assess chicken diversity at high genomic resolution. BMC genomics 20, 1–15

Maltecca, C., Tiezzi, F., Cole, J., and Baes, C. (2020). Symposium review: Exploiting homozygosity in the era of genomics—selection, inbreeding, and mating programs. Journal of dairy science 103, 5302–5313

Meuwissen, T. and Luo, Z. (1992). Computing inbreeding coefficients in large populations. Genetics Selection Evolution 24, 305–313

Meuwissen, T. H., Hayes, B. J., and Goddard, M. (2001). Prediction of total genetic value using genome-wide dense marker maps. genetics 157, 1819–1829

Meuwissen, T. H., Sonesson, A. K., Gebregiwergis, G., and Woolliams, J. A. (2020). Management of genetic diversity in the era of genomics. Frontiers in genetics 11, 880

Misztal, I., Lourenco, D., Aguilar, I., Legarra, A., and Vitezica, Z. (2018). Manual for BLUPF90 family of programs

Muir, W. (2007). Comparison of genomic and traditional blup-estimated breeding value accuracy and selection response under alternative trait and genomic parameters. Journal of Animal Breeding and Genetics 124, 342–355

Obsteter, J., Jenko, J., Hickey, J., and Gorjanc, G. (2019). Efficient use of genomic information for sustainable genetic improvement in small cattle populations. Journal of dairy science 102, 9971–9982

Perez-Enciso, M. (1995). Use of the uncertain relationship matrix to compute effective population size. Journal of Animal Breeding and Genetics 112, 327–332

Picard Druet, D., Varenne, A., Herry, F., Herault, F., Allais, S., Burlot, T., et al. (2020). Reliability of genomic evaluation for egg quality traits in layers. BMC genetics 21, 1–11

Pocrnic, I., Lourenco, D., Tsuruta, S., Chen, C., and Misztal, I. (2018). 327 practical problems and solutions using unknown parent groups in combined commercial pig sub-lines. Journal of Animal Science 96, 124

Pocrnic, I., Lourenco, D. A., Masuda, Y., and Misztal, I. (2016). Dimensionality of genomic information and performance of the algorithm for proven and young for different livestock species. Genetics Selection Evolution 48, 1–9

Qanbari, S., Hansen, M., Weigend, S., Preisinger, R., and Simianer, H. (2010). Linkage disequilibrium reveals different demographic history in egg laying chickens. BMC genetics 11, 1–10

Schaeffer, L. (2006). Strategy for applying genome-wide selection in dairy cattle. Journal of animal Breeding and genetics 123, 218–223

Sitzenstock, F., Ytournel, F., Sharifi, A. R., Cavero, D., Taubert, H., Preisinger, R., et al. (2013). Efficiency of genomic selection in an established commercial layer breeding program. Genetics Selection Evolution 45, 1–11

Sonesson, A. K., Woolliams, J. A., and Meuwissen, T. H. (2012). Genomic selection requires genomic control of inbreeding. Genetics Selection Evolution 44, 1–10

Vanavermaete, D., Fostier, J., Maenhout, S., and De Baets, B. (2020). Preservation of genetic variation in a breeding population for long-term genetic gain. G3: Genes, Genomes, Genetics 10, 2753–2762

VanRaden, P. M. (2008). Efficient methods to compute genomic predictions. Journal of dairy science 91, 4414–4423

Walsh, B. and Lynch, M. (2018). Evolution and selection of quantitative traits (Oxford University Press)

Waples, R. S. (2022). What is n e, anyway? Journal of Heredity 113, 371–379

Wientjes, Y. C., Bijma, P., Calus, M. P., Zwaan, B. J., Vitezica, Z. G., and van den Heuvel, J. (2022). The long-term effects of genomic selection: 1. response to selection, additive genetic variance, and genetic architecture. Genetics Selection Evolution 54, 1–21

Wiggans, G. R., Cole, J. B., Hubbard, S. M., and Sonstegard, T. S. (2017). Genomic selection in dairy cattle: the usda experience. Annual review of animal biosciences 5, 309–327

Wolc, A., Arango, J., Settar, P., Fulton, J. E., O’Sullivan, N. P., Preisinger, R., et al. (2011). Persistence of accuracy of genomic estimated breeding values over generations in layer chickens. Genetics Selection Evolution 43, 1–8

Wolc, A., Zhao, H. H., Arango, J., Settar, P., Fulton, J. E., O’sullivan, N. P., et al. (2015). Response and inbreeding from a genomic selection experiment in layer chickens. Genetics Selection Evolution 47, 1–12

Woolliams, J., Berg, P., Dagnachew, B., and Meuwissen, T. (2015). Genetic contributions and their optimization. Journal of Animal Breeding and Genetics 132, 89–99

Woolliams, J., Bijma, P., and Villanueva, B. (1999). Expected genetic contributions and their impact on gene flow and genetic gain. Genetics 153, 1009–1020

Wright, S. (1931). Evolution in mendelian populations. Genetics 16, 97

Zhang, J., Nie, C., Li, X., Ning, Z., Chen, Y., Jia, Y., et al. (2020). Genome-wide population genetic analysis of commercial, indigenous, game, and wild chickens using 600k snp microarray data. Frontiers in genetics 11, 543294

