## Supplemental Files for "Assessment of long-term trends in genetic mean and variance after the introduction of genomic selection in layers: a simulation study"

### Supplementary Material

#### 1 SUPPLEMENTARY TABLES

##### 1.1 Supplementary Table 1

**Table S1.** Rate of inbreeding ($\Delta F$ x100) and effective population size ($Ne$) based on observed neutral loci heterozygosity (SD x100 over replicates in parentheses).

| Breeding program | 40 sires per generation | | | 120 sires per generation | | |
|---|---|---|---|---|---|---|
| | $\Delta F$/year (x100) | $\Delta F$/generation (x100) | $Ne$ | $\Delta F$/year (x100) | $\Delta F$/generation (x100) | $Ne$ |
| PTS | 5.15 (0.49) | 5.15 (0.49) | 10 | 3.19 (0.20) | 3.19 (0.20) | 16 |
| GTS | 4.81 (0.14) | 3.60 (0.14) | 14 | 3.60 (0.35) | 2.70 (0.26) | 19 |
| GTSMF | 4.67 (0.18) | 3.50 (0.20) | 14 | 3.61 (0.24) | 2.71 (0.18) | 19 |
| GOCS 45 | 3.54 (0.18) | 2.66 (0.13) | 19 | 2.89 (0.17) | 2.16 (0.13) | 23 |
| GOCS 65 | 2.23 (0.05) | 1.67 (0.03) | 30 | 1.74 (0.16) | 1.31 (0.12) | 38 |

| | Fluctuating number of sires per generation | | |
|---|---|---|---|
| | $\Delta F$/year (x100) | $\Delta F$/generation (x100) | $Ne$ |
| UGOCS 45 | 5.50 (0.39) | 4.15 (0.31) | 12 |
| UGOCS 55 | 3.24 (0.16) | 2.44 (0.12) | 21 |

*PTS - conventional truncation selection; GTS - genomic truncation selection;*
*GTSMF - GTS with minimization of progeny inbreeding;*
*GOCS X - genomic optimal contribution selection; UGOCS X - unconstrained GOCS; with the X trigonometric pe*

#### 1.2 Supplementary Table 2

**Table S2.** Rate of inbreeding ($\Delta F$ x100) and effective population size ($Ne$) based on observed QTL heterozygosity (SD x100 over replicates in parentheses).

| Breeding program | 40 sires per generation | | | 120 sires per generation | | |
|---|---|---|---|---|---|---|
| | $\Delta F$/year (x100) | $\Delta F$/generation (x100) | $Ne$ | $\Delta F$/year (x100) | $\Delta F$/generation (x100) | $Ne$ |
| PTS | 5.27 (0.44) | 5.27 (0.44) | 9 | 3.26 (0.22) | 3.26 (0.22) | 15 |
| GTS | 4.95 (0.16) | 3.71 (0.12) | 13 | 3.74 (0.36) | 2.81 (0.27) | 18 |
| GTSMF | 4.97 (0.23) | 3.73 (0.17) | 13 | 3.76 (0.25) | 2.82 (0.19) | 18 |
| GOCS 45 | 3.70 (0.24) | 2.77 (0.18) | 18 | 2.96 (0.16) | 2.22 (0.12) | 22 |
| GOCS 65 | 2.35 (0.06) | 1.77 (0.05) | 28 | 1.79 (0.15) | 1.35 (0.11) | 37 |
| | Fluctuating number of sires per generation | | | | | |
| | $\Delta F$/year (x100) | $\Delta F$/generation (x100) | $Ne$ | | | |
| UGOCS 45 | 5.68 (0.39) | 4.26 (0.27) | 12 | | | |
| UGOCS 55 | 3.35 (0.15) | 2.51 (0.11) | 20 | | | |

*PTS - conventional truncation selection; GTS - genomic truncation selection;*
*GTSMF - GTS with minimization of progeny inbreeding;*
*GOCS X - genomic optimal contribution selection; UGOCS X - unconstrained GOCS; with the X trigonometric p*

#### 1.3 Supplementary Table 3

**Table S3.** Rate of inbreeding ($\Delta F$ x100) and effective population size ($Ne$) based on pedigree (SD x100 over replicates in parentheses).

| Breeding program | 40 sires per generation | | | 120 sires per generation | | |
|---|---|---|---|---|---|---|
| | $\Delta F$/year (x100) | $\Delta F$/generation (x100) | $Ne$ | $\Delta F$/year (x100) | $\Delta F$/generation (x100) | $Ne$ |
| PTS | 3.35 (0.52) | 3.35 (0.52) | 20 | 1.47 (0.15) | 1.47 (0.15) | 36 |
| GTS | 0.93 (0.06) | 0.70 (0.04) | 72 | 0.34 (0.01) | 0.25 (0.01) | 196 |
| GTSMF | 0.85 (0.09) | 0.64 (0.07) | 78 | 0.33 (0.03) | 0.25 (0.02) | 202 |
| GOCS 45 | 0.25 (0.02) | 0.19 (0.01) | 267 | 0.11 (0.01) | 0.09 (0.01) | 584 |
| GOCS 65 | 0.17 (0.01) | 0.13 (0.01) | 390 | 0.07 (0.01) | 0.07 (0.01) | 994 |

| | Fluctuating number of sires per generation | | |
|---|---|---|---|
| | $\Delta F$/year (x100) | $\Delta F$/generation (x100) | $Ne$ |
| UGOCS 45 | 1.14 (0.07) | 0.85 (0.05) | 59 |
| UGOCS 55 | 0.25 (0.01) | 0.19 (0.01) | 267 |

*PTS - conventional truncation selection; GTS - genomic truncation selection;*
*GTSMF - GTS with minimization of progeny inbreeding;*
*GOCS X - genomic optimal contribution selection; UGOCS X - unconstrained GOCS; with the X trigonometric pe*

### 2 SUPPLEMENTARY FIGURES

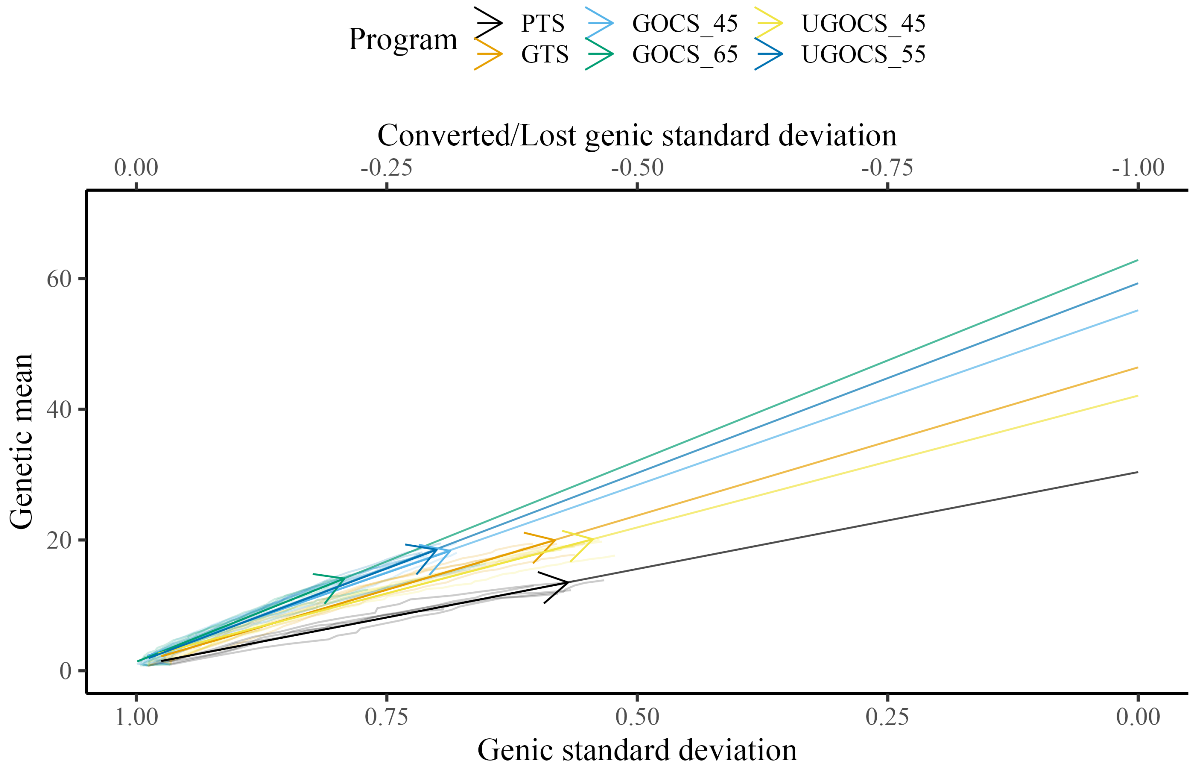


**Figure 1a.**

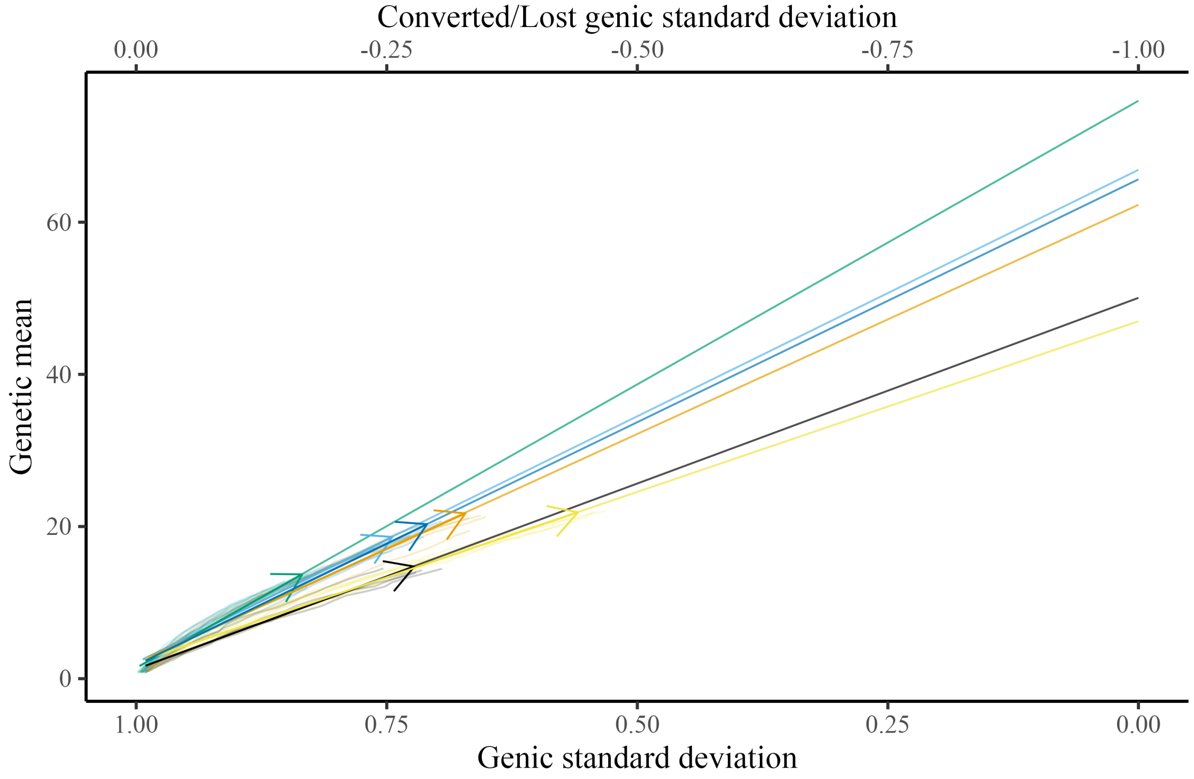


**Figure 1b.**

**Figure 1.** Conversion efficiency for conventional truncation selection (PTS) program and genomic programs (genomic truncation selection - GTS, genomic optimal contribution selection - GOCS X, unconstrained GOCS - UGOCS X, with the X trigonometric penalty degrees) marked with an arrow and further extrapolated to 100% of genic variance lost for **(A)** 40 sires and **(B)** 120 sires scenario.

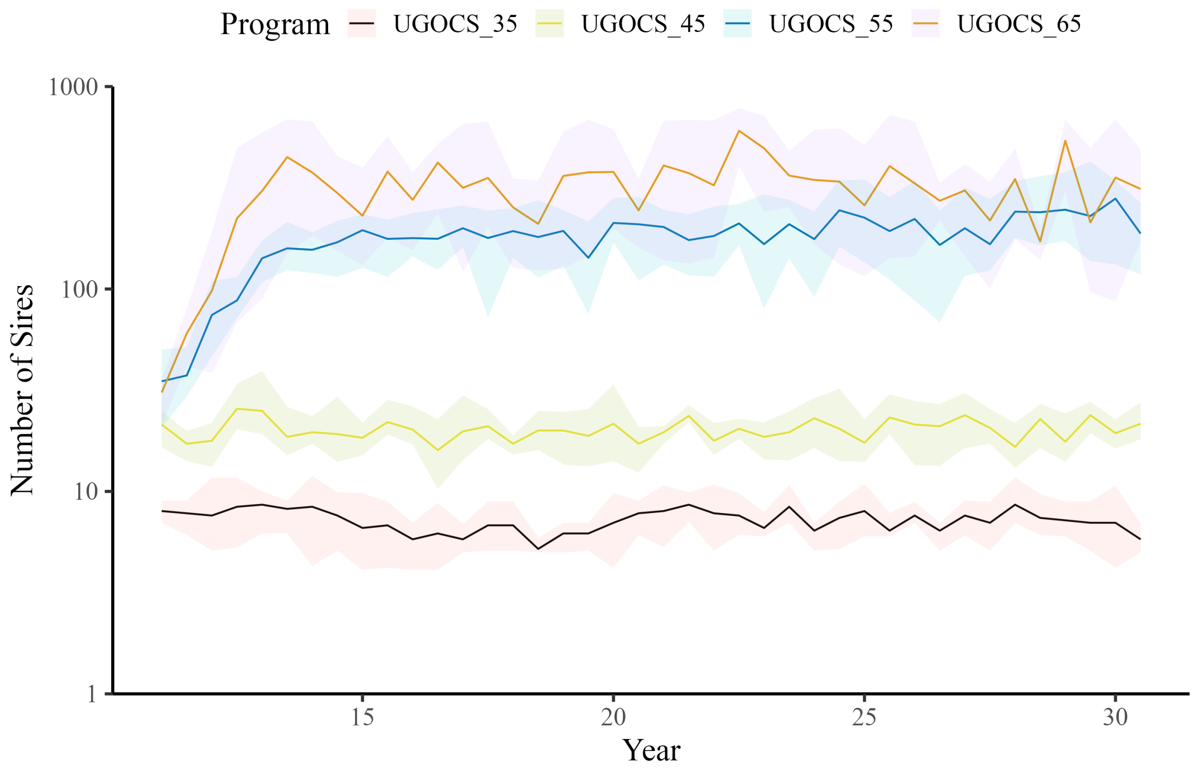


**Figure 2a.**

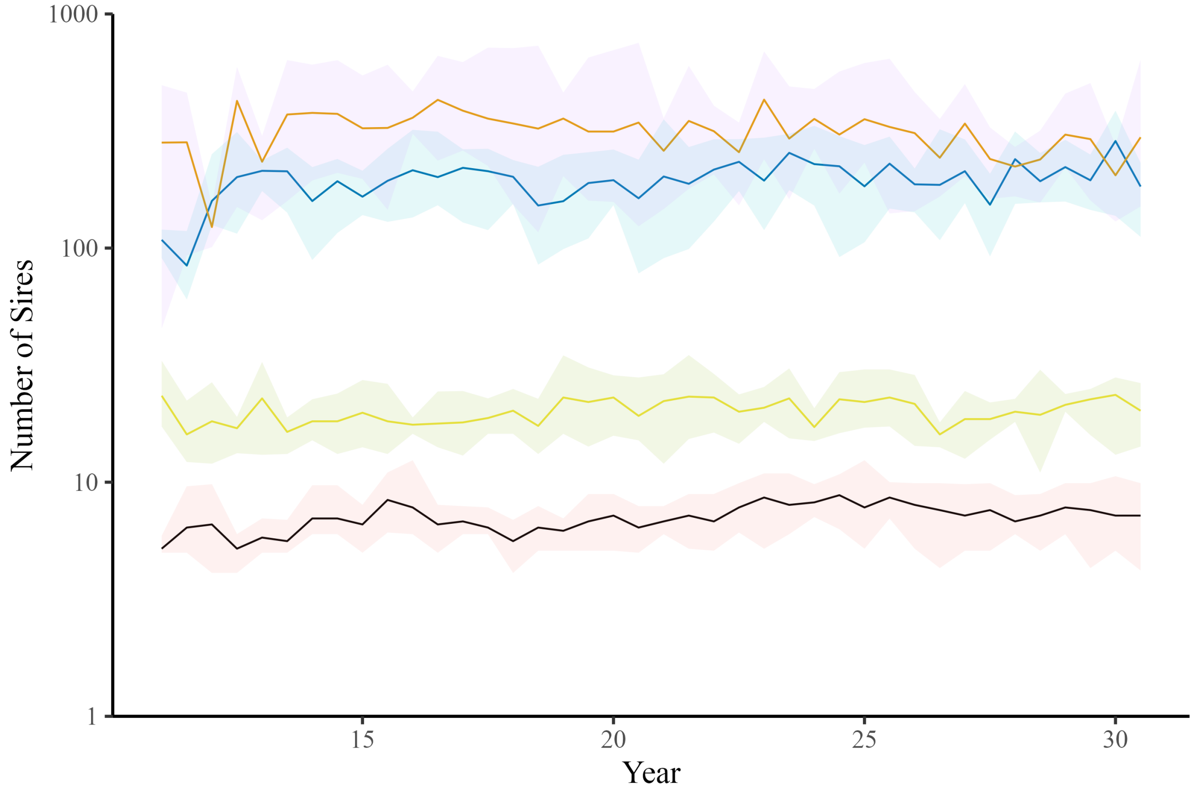


**Figure 2b.**

**Figure 2.** Number of sires in each year of genomic unconstrained optimal contribution selection (UGOCS) programs for **(A)** 40 sires and **(B)** 120 sires scenario.

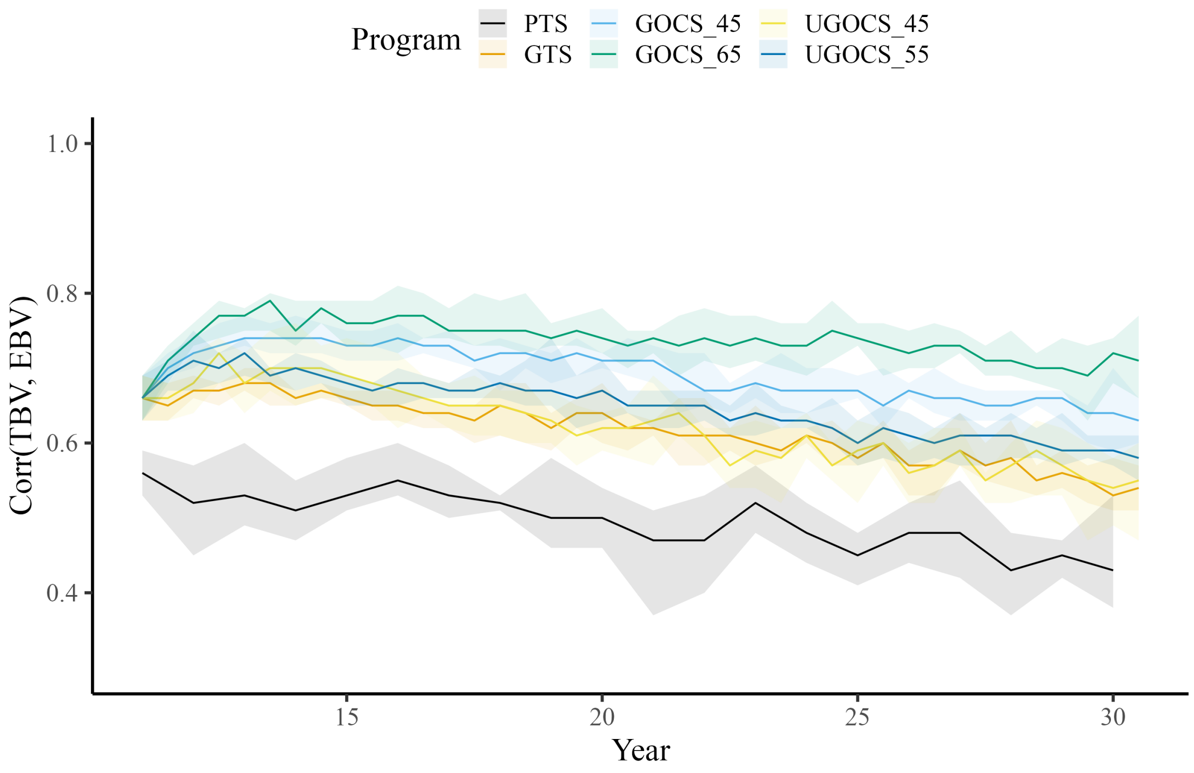


**Figure 3a.**

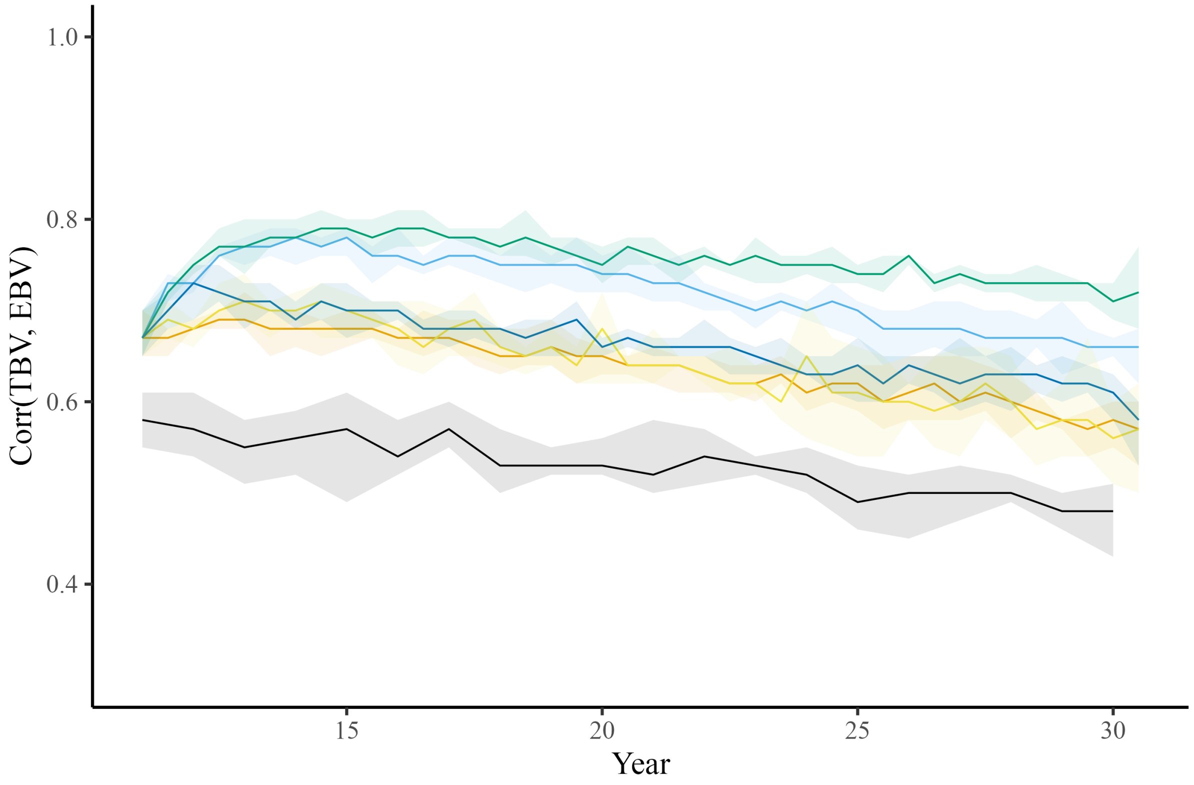


**Figure 3b.**

**Figure 3.** Accuracy of selection candidates for conventional truncation selection (PTS) program and genomic programs (genomic truncation selection - GTS, genomic optimal contribution selection - GOCS X, unconstrained GOCS - UGOCS X, with the X trigonometric penalty degrees) across the 20 years of selection for **(A)** 40 sires and **(B)** 120 sires scenario.

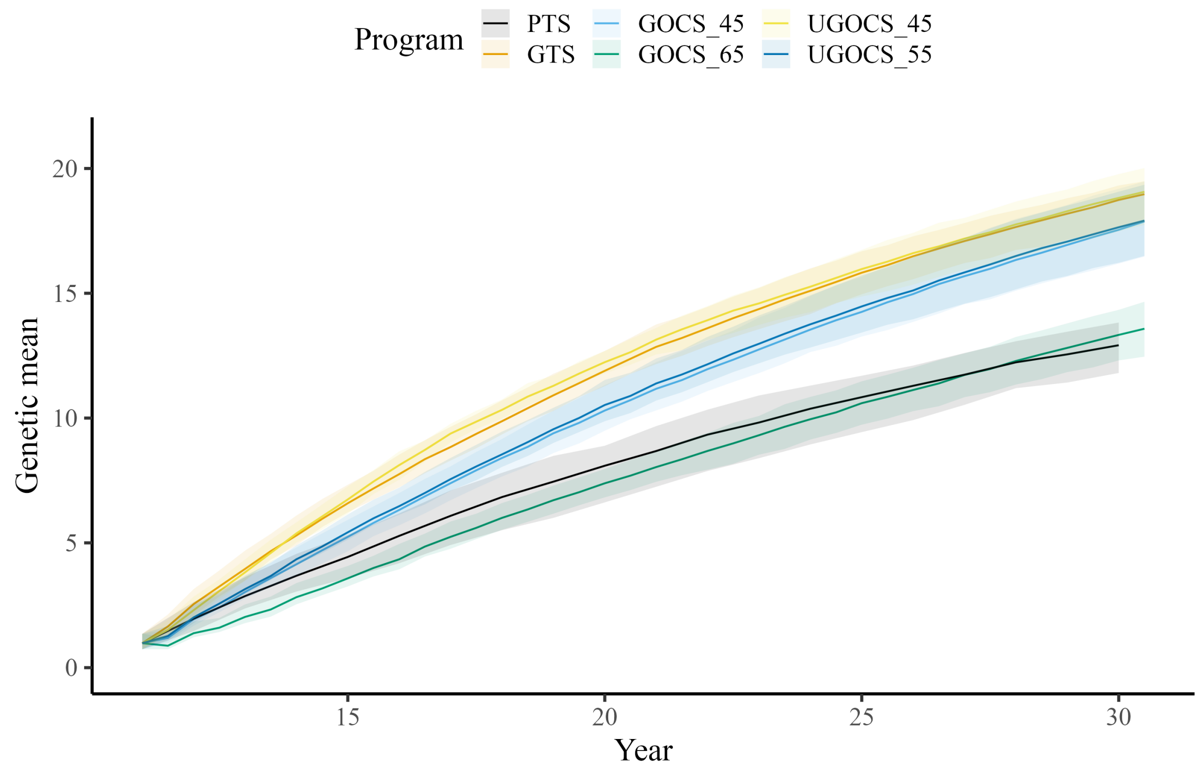


**Figure 4a.**

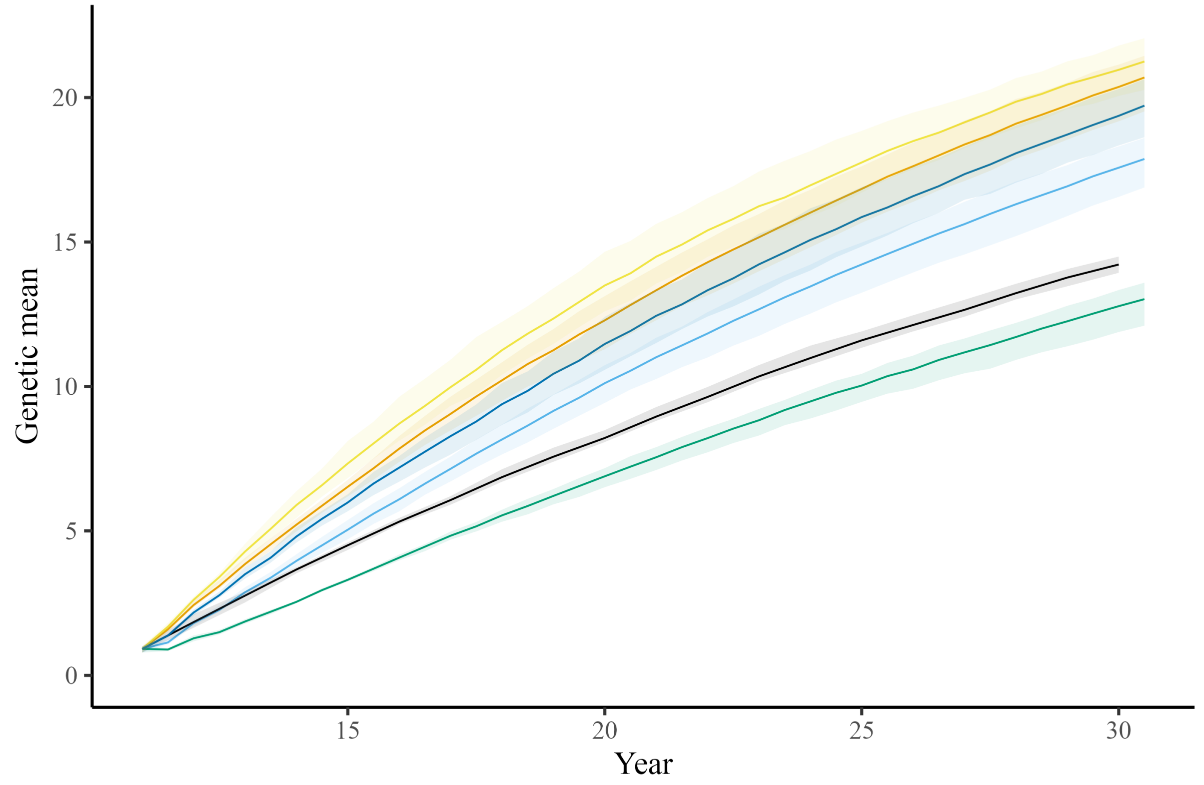


**Figure 4b.**

**Figure 4.** Genetic mean of selection candidates for conventional truncation selection (PTS) program and genomic programs (genomic truncation selection - GTS, genomic optimal contribution selection - GOCS X, unconstrained GOCS - UGOCS X, with the X trigonometric penalty degrees) across the 20 years of selection for **(A)** 40 sires and **(B)** 120 sires scenario.

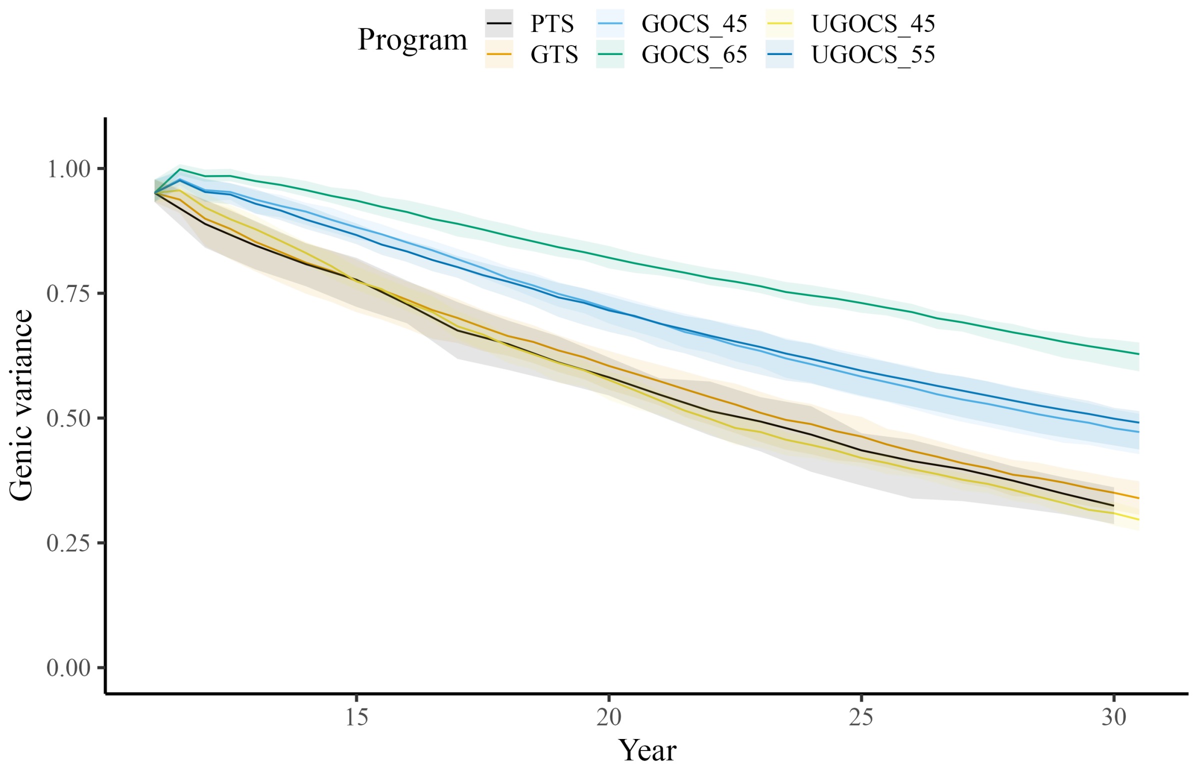


**Figure 5a.**

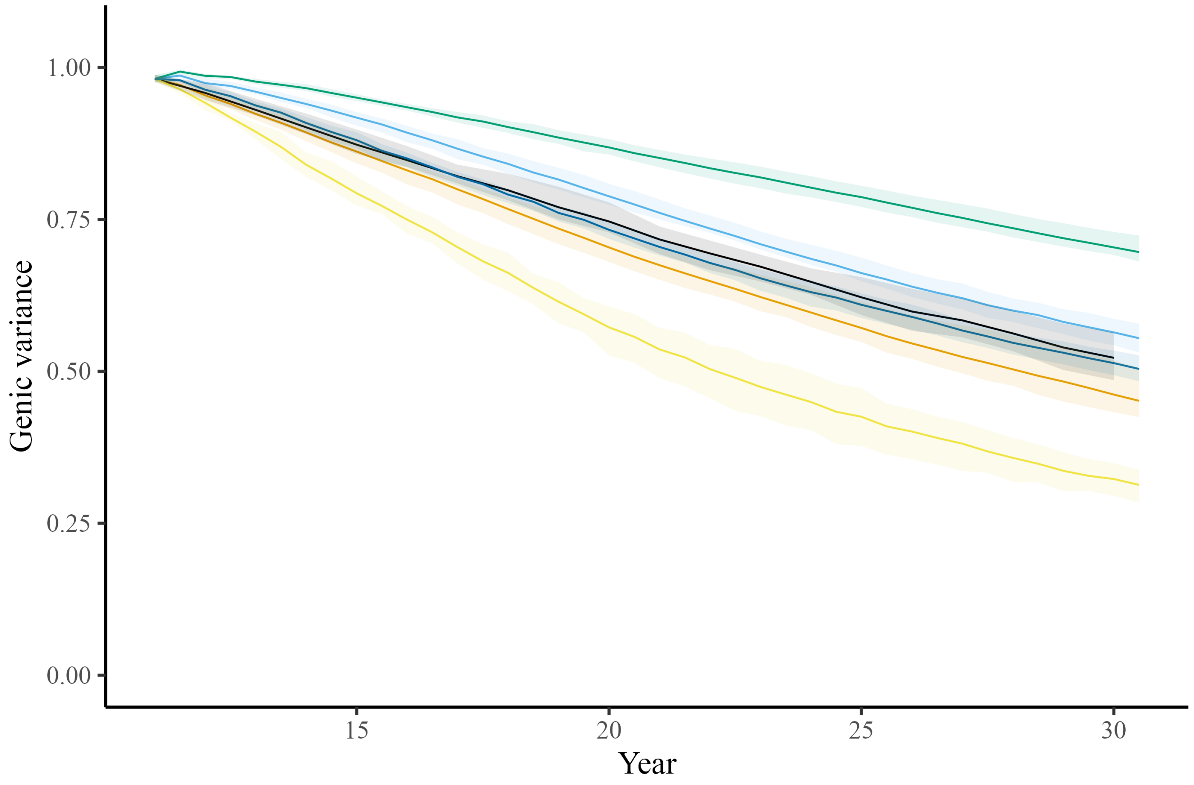


**Figure 5b.**

**Figure 5.** Genic variance of selection candidates for conventional truncation selection (PTS) program and genomic programs (genomic truncation selection - GTS, genomic optimal contribution selection - GOCS X, unconstrained GOCS - UGOCS X, with the X trigonometric penalty degrees) across the 20 years of selection for **(A)** 40 sires and **(B)** 120 sires scenario.
